# Structure-Guided Discovery and Characterization of Novel FLT3 Inhibitors for Acute Myeloid Leukemia Treatment

**DOI:** 10.1101/2025.06.26.661886

**Authors:** Bishal Budha, Gourab Basu Choudhury, Md. Shohag Hossain, Arjun Acharya

## Abstract

FLT3 (FMS-like tyrosine kinase 3), a receptor tyrosine kinase, is frequently mutated in acute myeloid leukemia (AML), a hematologic malignancy marked by aggressive proliferation, poor prognosis, and high relapse rates. Although FDA-approved FLT3 inhibitors exist, their clinical efficacy is often undermined by resistance and off-target effects, underscoring the critical necessity for more effective and selective agents. Here, we employed a structure-based computational approach combining pharmacophore screening via Pharmit and the MolPort compound library to identify novel FLT3 inhibitors. Pharmacophore modeling, virtual screening, and docking identified two promising leads, MolPort-002-705-878 and MolPort-007-550-904, with binding affinities of −11.33 and −10.66 kcal/mol, correspondingly. These compounds were further characterized through molecular dynamics (MD) simulations, incorporating Principal Component Analysis (PCA), free energy landscape (FEL) analysis, density functional theory (DFT) calculations, and ADMET profiling. MD results supported the integrity of the FLT3–lead complexes; DFT revealed favorable reactivity, and ADMET predictions indicated drug-likeness with low toxicity, pending experimental confirmation. This integrated *in silico* pipeline highlights the potential of these molecules as next-generation FLT3 inhibitors and offers a scalable strategy for targeted AML therapeutics.

## 1 Introduction

Acute myeloid leukemia (AML) is a fast-progressing blood cancer marked by the unchecked proliferation of immature myeloid cells in the bone marrow and bloodstream [1]. This clonal expansion disrupts normal blood cell production, resulting in bone marrow failure and swift decline in patient health [1, 2]. Representing the most recurrent type of acute leukemia in mature individuals, AML accounts for over half of all adult leukemia cases and is associated with particularly poor outcomes in high-risk and elderly populations, where five-year survival rates remain below 30% [3, 4]. Despite advances in identifying key genetic mutations such as FLT3, IDH1/2, and NPM1, and the curative potential of bone marrow transplantation in select patients, acute myeloid leukemia (AML) remains largely incurable, particularly in cases of relapse or treatment resistance. Moreover, AML is known for its rapid progression , and if left untreated, it can become increasingly aggressive, further emphasizing the urgent need for more effective and durable therapeutic strategies [5–7].

FMS-like tyrosine kinase 3 (FLT3) is a receptor tyrosine kinase predominantly expressed in hematopoietic stem cells and plays a pivotal role in the regulation of hematopoiesis [5]. Mutations in FLT3, particularly internal tandem duplications (FLT3-ITD) and tyrosine kinase domain point mutations (FLT3-TKD), are among the most frequent genetic abnormalities in acute myeloid leukemia (AML), affecting nearly 30% of patients [8, 9]. As a result of these mutations, the kinase remains persistently active, which promotes malignant signaling and is associated with adverse prognosis [10]. Consequently, FLT3 has gained recognition as a proven biological site in the therapeutic management of AML.

FLT3 inhibitors have established as promising therapeutic agents for AML, supported by a wide array of preclinical and clinical studies [11–13]. Agents like Midostaurin, Quizartinib, and Gilteritinib exert their activity by targeting the binding cavity for ATP within FLT3 receptor, thereby preventing its phosphorylation and downstream signaling [13–15]. Although all of these inhibitors have gained regulatory approval and are commercially available in the United States, their clinical effectiveness is often compromised by issues such as acquired resistance, off-target effects, and suboptimal specificity [14, 16]. These limitations not only reduce therapeutic efficacy but also affect patient safety and quality of life. As a result, there remains a pressing need to design and develop next-generation FLT3 inhibitors with enhanced potency, selectivity, and pharmacological profiles to achieve better clinical outcomes in AML therapy.

Structure-Based Virtual Screening (SBVS) is a key component of modern CADD (Computer-Aided Drug Design), enabling the identification of candidate molecules based on their structural compatibility with the target binding site [17]. Among available platforms, Pharmit offers distinct advantages through its interactive, cloud-based interface that integrates pharmacophore modeling, shape screening, and energy minimization [18]. It allows real-time refinement of pharmacophore queries directly from protein-ligand complexes and supports rapid filtering of large compound libraries based on steric and electronic features [18].

Molecular docking estimates the most favorable binding pose and interaction strength of ligands at receptor sites, making it a key component of structure-based drug design [19]. However, its static nature limits understanding of dynamic interactions. To address this, molecular dynamics (MD) simulations are carried out to examine temporal stability, conformational mobility, and ligand persistence under near-physiological conditions [20]. Post-MD MM/GBSA calculations refine hit selection by quantifying binding free energies, accounting for solvation and entropic contributions [21]. Density Functional Theory (DFT) provides deep insights into electronic properties such as frontier orbitals and charge distribution, aiding in reactivity and binding specificity analysis [22]. Finally, *in silico* ADMET profiling evaluates pharmacokinetics and toxicity, ensuring drug-likeness and safety of the candidate compounds [23].

This study presents an integrated computational strategy comprising pharmacophore modeling (via the Pharmit platform), molecular docking, MD simulations with PCA and FEL analysis, DFT calculations, and ADMET evaluation to guide the rational design of FLT3 inhibitors. By combining these complementary methods, the approach aims to systematically identify and characterize hit compounds with strong structural compatibility, binding stability, favorable electronic properties, and suitable pharmacokinetics. The eventual objective is to discover novel chemical scaffolds with strong potential to inhibit FLT3, advancing targeted therapies for acute myeloid leukemia (AML).

## 2 Materials and methods

### Pharmacophore Modeling and Virtual Screening

In this study, we examined the crystallographic conformation of the FLT3 kinase complexed with the FDA-approved drug, Gilteritinib (PDB ID: 6JQR) [24], using it as a reference to identify key molecular interactions and define relevant pharmacophoric features. A ligand-based pharmacophore model was subsequently developed from this FLT3 crystal structure using the Pharmit web server [18].

This pharmacophore was employed to screen the MolPort compound library, one of the most recently curated and expansive repositories, containing 4,742,020 compounds as of September 5, 2024. All candidates were filtered according to Lipinski’s Rule of Five [25] for ensuring drug-likeness. Screening was performed with only the best conformer per compound to maintain computational efficiency. From the resulting dataset, the top 25,000 hits were selected, followed by energy minimization within the server to refine ligand geometries. Compounds exhibiting a maximum minimized binding affinity (docking score) of zero were retained. Finally, a threshold value of 1Åfor molecular root-mean-square deviation (mRMSD) was enforced, and compounds showing substantial deviation from the pharmacophore alignment were excluded, ensuring high fidelity between the screened conformations and the original pharmacophore model.

### Preprocessing of Ligands and Protein

The X-ray crystallographic structure of the FLT3 receptor (PDB ID: 6JQR) [24] was retrieved from the RCSB Protein Data Bank [26]. Homology modeling was performed using MODELLER to fill in missing residues and generate a structurally complete protein model [27]. Protein preparation was conducted using AutoDockTools [28], during which all water molecules were deleted, polar hydrogens were introduced, and Kollman partial charges were applied. Three-dimensional structures of the identified hits from pharmacophore-based virtual screening were imported from the screening server. Following energy minimization and charge assignment, PDB format files of both protein and ligands were converted into PDBQT format using OpenBabel [29].

### Molecular Docking

The receptor grid for docking was constructed around key catalytic residues of FLT3; Leu616, Val624, Ala642, Glu692, Cys694, Asp698, and Leu818 based on their involvement in the active site [30], using grid dimensions of 20Å, 25Å, and 22Åwith a grid point spacing of 1Åalong the *x*, *y*, and *z* axes, respectively. Docking-based virtual screening was performed within that receptor grid using PyRx where autodock vina serves in backend [29]. The cognate ligand was redocked into its binding site, and the RMSD between its crystallographic and predicted poses was calculated to validate the docking procedure. The resulting top complexes of protein-ligand were visually examined using PyMOL 2.5.2 [31], while the molecular interactions were further analyzed using BIOVIA Discovery Studio Visualizer [32].

### Molecular Dynamics Simulations

GROMACS 2025.1 [33] was employed to conduct molecular dynamics simulations aimed at probing the structural stability of the protein and its ligand-bound forms [34]. The protein was parametrized deploying CHARMM General Force Field (CGenFF), and ligand topologies were generated via SwissParam [35]. Structures underwent energy minimization for 2500 steps using the steepest descent algorithm. The systems were solvated using the TIP3P water model [36] and neutralized with Na^+^ and Cl^−^ ions using `gmx genion`. Two equilibration phases followed: a 100-ps NVT run to reach 300 K, and a 100-ps NPT run for pressure and density stabilization. During equilibration, all bond lengths were constrained to preserve structural integrity, and water constraints facilitated solvent shell reorganization. The v-rescale thermostat and Parrinello–Rahman barostat [37] were employed for temperature and pressure control, respectively. The LINCS algorithm ensured bond constraints [38], and long-range electrostatics were calculated using the PME method [39]. Production MD simulations were executed for 100 ns to trace trajectory and stability analysis.

### MM/GBSA Binding Free Energy

Binding free energies were calculated using the Molecular Mechanics/Generalized Born Surface Area (MM/GBSA) method via the `gmx MMPBSA` tool (version 1.6.3), which integrates GROMACS with AmberTools for end-state energy analysis [40]. The overall binding free energy (Δ*G*_bind_) was partitioned into van der Waals (Δ*E*_vdW_), electrostatic (Δ*E*_ele_), and both polar and non-polar solvation energy contributions, as outlined in Equation 1 [34].

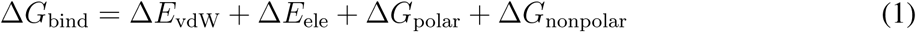

Polar solvation energy was evaluated by the Generalized Born (GB) using the GB-Neck2 implicit solvent model (`igb=5`), and the nonpolar term was estimated via the LCPO method [41]. Entropy contributions were omitted, consistent with common practice in comparative MM/GBSA studies [42]. A total of 1001 snapshots were extracted at equal intervals from the 100 ns production trajectory (`MD center.xtc`) for free energy analysis.

### Principal Component Analysis (PCA)

Trajectory pre-processing and PCA execution were performed using `GROMACS 2025.1` [33]. The positional atomic covariance matrix was computed using the `gmx covar` command, focusing on the C*α* atoms of the protein to capture backbone motions. This step produced eigenvectors along with their corresponding eigenvalues. Subsequently, the top three principal components were then analyzed using the `gmx anaeig` utility to examine root-mean-square (RMS) fluctuations, eigenvector contributions, and 2D trajectory projections. All PCA plots were generated using `Xmgrace` [43].

### Free Energy Landscape (FEL)

The trajectory file from the 100 ns molecular dynamics simulation was taken as input for Principal Component Analysis (PCA), from which the top two eigenvectors (PC1 and PC2) were selected as reaction coordinates. The free energy of each conformation was calculated on the basis of free energy Δ*G*, Boltzmann constant (*k_B_* = 1.380649 *×* 10^−23^ J/K), absolute temperature (T=298.15 K) and natural logarithm of the normalized probability distribution across the PC1–PC2 space (ln *P* ), using the Boltzmann relation as shown in Equation 2 [44].

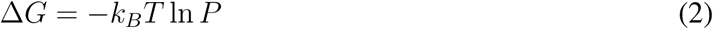

The three-dimensional FEL was visualized using the matplotlib.pyplot.plot surface() function to map local energy minima and transition states on a rugged conformational landscape [45].

### DFT Analysis

The molecular structure and Gaussian input files of the leads were generated with the GaussView 6 program [46]. The molecular geometries of the leads were fully optimized in the gas phase employing density functional theory at the B3LYP level and the 6-311G(d,p) basis set, facilitated by the Gaussian 09W software package [47]. Optimized molecular structures were used to compute various molecular properties, including frontier molecular orbitals (HOMO & LUMO), the energy gaps (Δ E), quantum chemical descriptors, molecular electrostatic potential (MEP) map, Mulliken charges and natural bond orbital (NBO) study. Key global reactivity descriptors such as ionization potential (*I*), electron affinity (*A*), chemical hardness (*η*), chemical softness (*S*), chemical potential (*µ*), global electrophilicity index (*ω*) and, Nucleophilicity index (N) were derived from orbital energy values on the basis of Koopman’s principles (Equations 3 & 4) [48, 49].

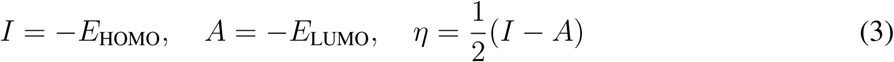

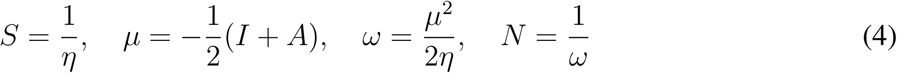

The NBO program [50] was employed to investigate natural bond orbitals at the same level of theory, where delocalized interactions were analyzed using second-order perturbation theory. The stabilization energy *E*(2) was computed from the donor orbital occupancy (*q_i_*), donor and acceptor energies (*E_i_, E_j_*), and the Fock matrix off-diagonal element (*F_ij_*) as per Equation 5 [51, 52].

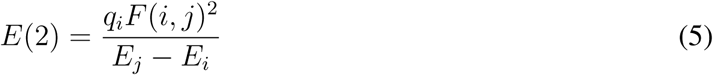

### ADMET Analysis

The compounds with the most favorable docking scores were further evaluated for their pharmacokinetic properties using the SwissADME web server [23], and for toxicity using the ProTox-3.0 web server [53].

## 3 Results and discussion

### Pharmacophore-based Virtual Screening

The pharmacophore query model constructed using the key interaction features of Gilteritinib observed within the FLT3 binding site (Figure 2 and 1S) incorporated four critical pharmacophoric features; one hydrophobic region, one hydrogen bond acceptor, and two hydrogen bond donor groups, each of which was deemed essential for a compound to be considered a valid hit. The precise spatial coordinates (X, Y, and Z) of these features are detailed in Table 1. Utilizing this defined feature set, a ligand-based virtual screening was performed against the MolPort compound library and only those molecules matching all four features were retained. From the initial top 25,000 compounds identified as potential hits by the server, 439 compounds (Table 1S) met the screening criteria: compliance with Lipinski’s Rule of Five, minimized binding affinity less than zero, and a molecular RMSD (mRMSD) below 1Å. These 439 hits were selected (Table 2S) and transferred for further validation and refinement through docking-based virtual screening.

**Figure 1:**
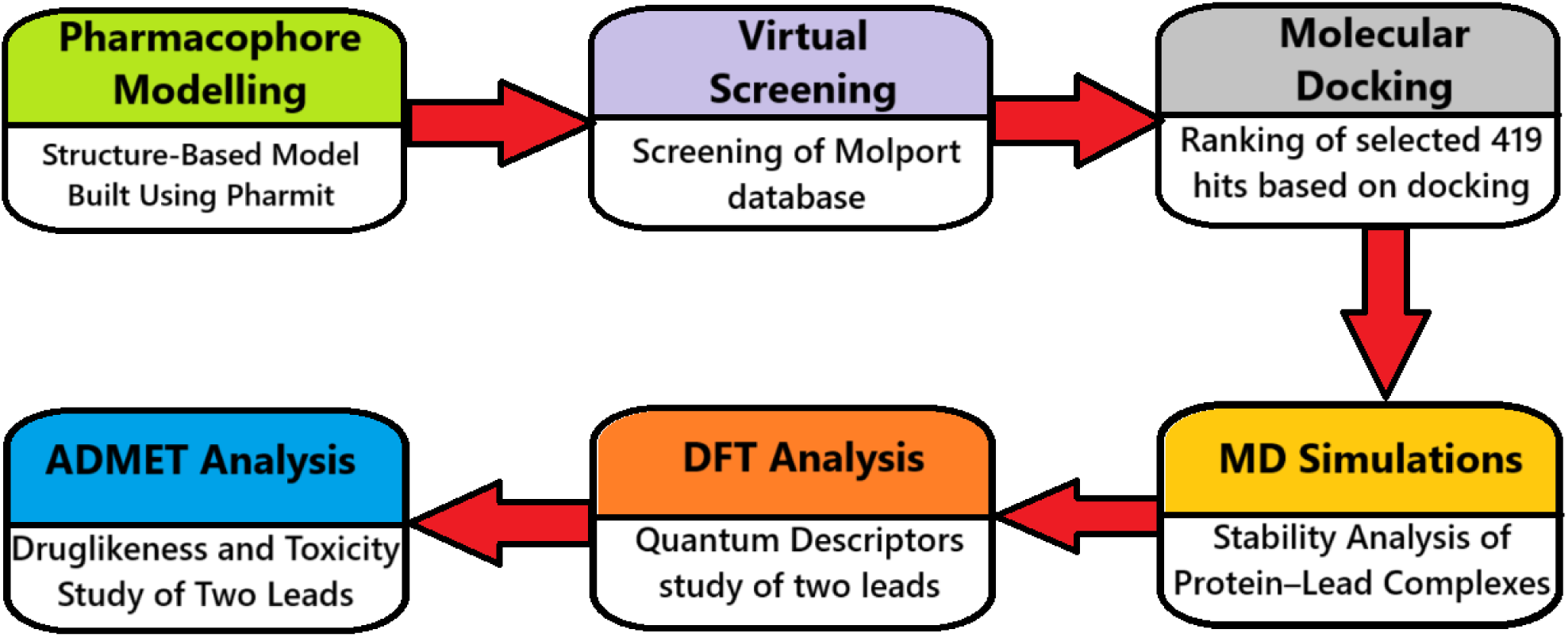
Workflow diagram providing a concise overview of the key steps and structure of the study.

**Figure 2:**
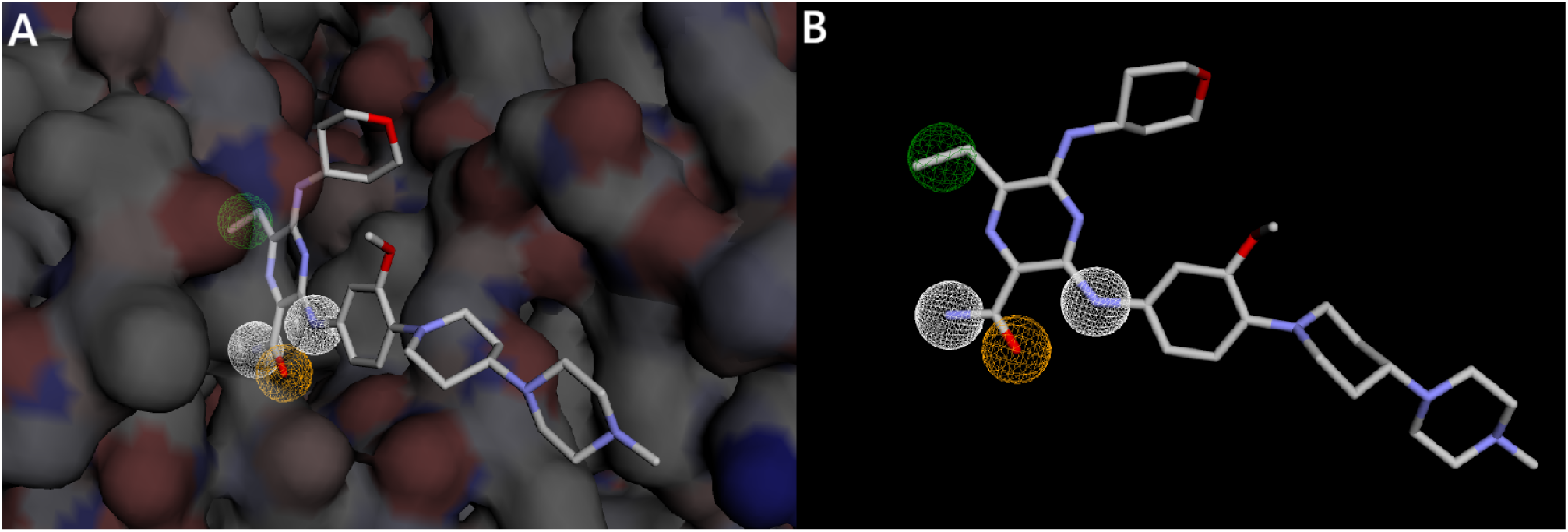
Pharmacophore features generated by Pharmit server; (A) within structure (B) ligand-basis. Two white zone represent hydrogen donor region, orange represents Hydrogen Acceptor while green represents Hydrophobic.

**Table 1:** Pharmacophore features and computed 3D Coordinates of their from Pharmit.

| Parameter | Hydrogen Donor(1) | Hydrogen Donor(2) | Hydrogen Acceptor | Hydrophobic |
| --- | --- | --- | --- | --- |
| X | -28.282 | -27.437 | -27.632 | -28.146 |
| Y | -7.483 | -3.380 | -5.511 | -2.242 |
| Z | -30.381 | -31.714 | -32.244 | -27.314 |
| Radius | 1 | 1 | 1 | 1 |

**Table 2:**
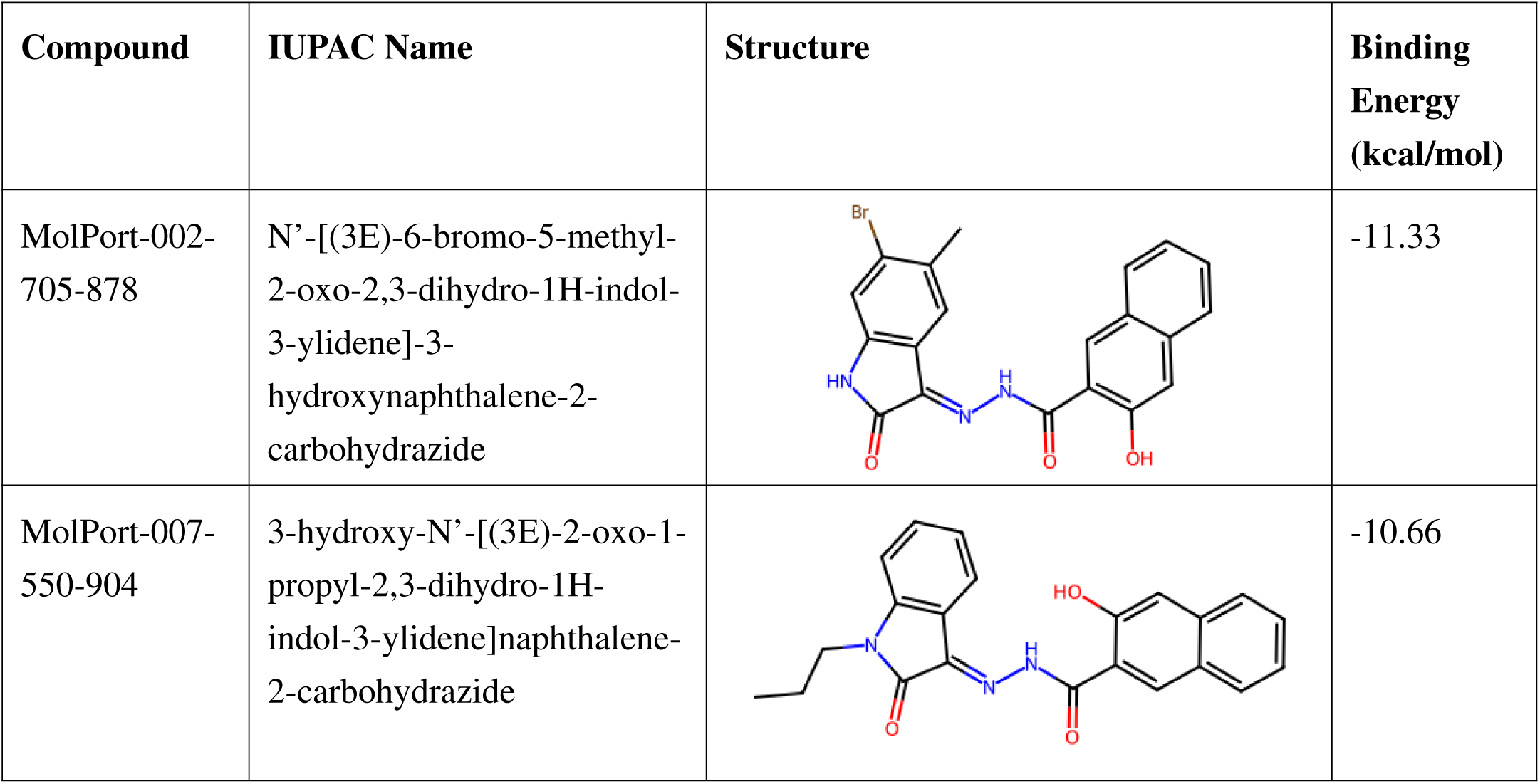
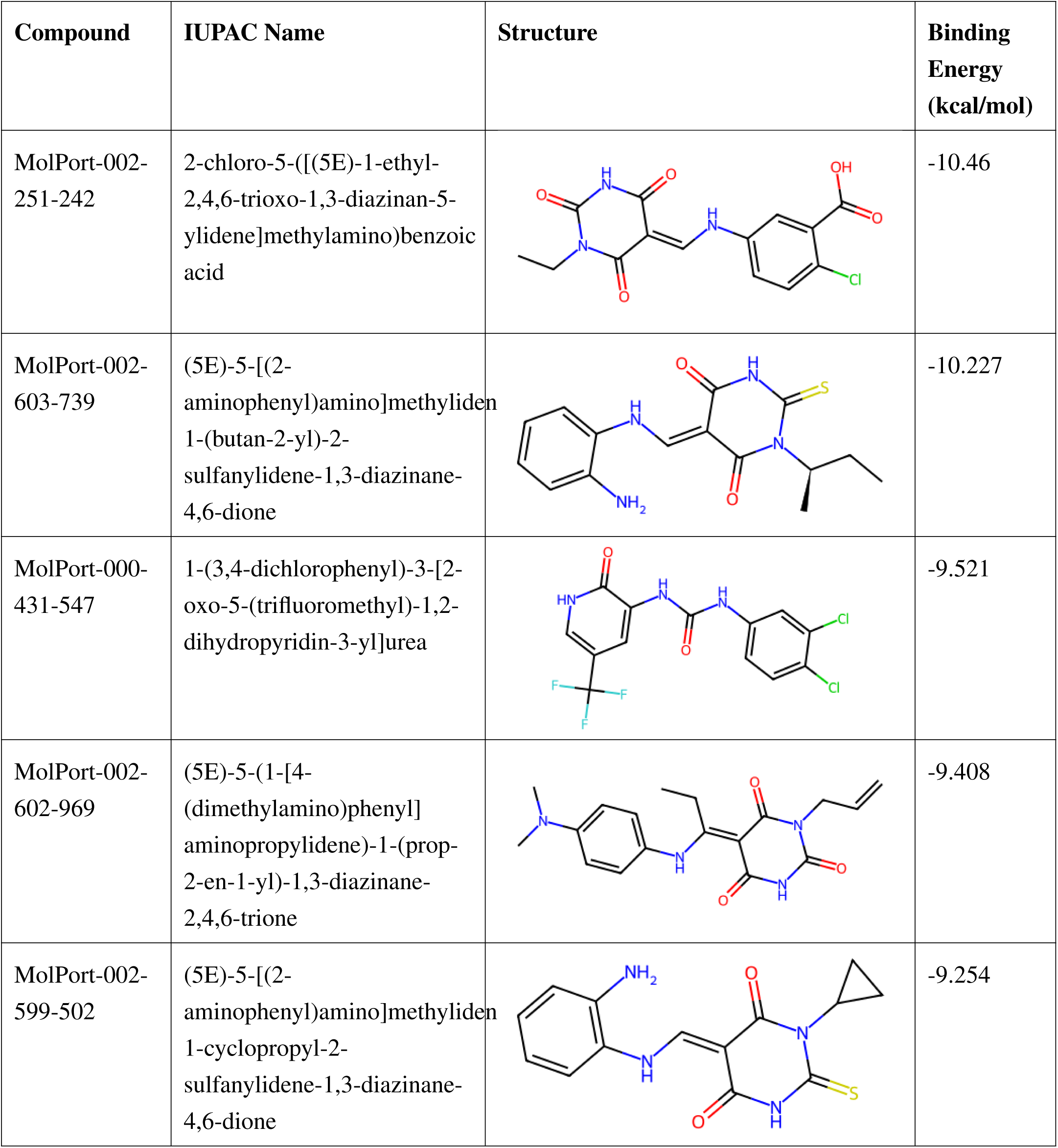
Binding energies and structures of selected compounds.

| Compound | IUPAC Name | Structure | Binding Energy (kcal/mol) |
| --- | --- | --- | --- |
| MolPort-002-705-878 | N'-[(3E)-6-bromo-5-methyl-2-oxo-2,3-dihydro-1H-indol-3-ylidene]-3-hydroxynaphthalene-2-carbohydrazide |  | -11.33 |
| MolPort-007-550-904 | 3-hydroxy-N'-[(3E)-2-oxo-1-propyl-2,3-dihydro-1H-indol-3-ylidene]naphthalene-2-carbohydrazide |  | -10.66 |
| MolPort-002-251-242 | 2-chloro-5-([(5E)-1-ethyl-2,4,6-trioxo-1,3-diazinan-5-ylidene]methylamino)benzoic acid |  | -10.46 |
| MolPort-002-603-739 | (5E)-5-[(2-aminophenyl)amino]methyliden-1-(butan-2-yl)-2-sulfanylidene-1,3-diazinane-4,6-dione |  | -10.227 |
| MolPort-000-431-547 | 1-(3,4-dichlorophenyl)-3-[2-oxo-5-(trifluoromethyl)-1,2-dihydropyridin-3-yl]urea |  | -9.521 |
| MolPort-002-602-969 | (5E)-5-(1-[4-(dimethylamino)phenyl]aminopropylidene)-1-(prop-2-en-1-yl)-1,3-diazinane-2,4,6-trione |  | -9.408 |
| MolPort-002-599-502 | (5E)-5-[(2-aminophenyl)amino]methyliden-1-cyclopropyl-2-sulfanylidene-1,3-diazinane-4,6-dione |  | -9.254 |

### Docking based virtual screening

The total of 419 hits from pharmacophore modeling were subjected to molecular docking with FLT3 within the defined receptor grid. The resulting docking scores spanned from —11.330 to –3.052 kcal/mol. Based on these results, the top seven highest-ranking compounds crossing binding energy below –9 kcal/mol were shortlisted with their RDKit [54] generated structure image and IUPAC name in Table 2. The binding affinity profiles of these ligands suggest a significant potential to suppress the functional activity of the target protein.

Docking validation was performed by re-docking the cognate ligand Gilteritinib into its native binding zone. The resulting RMSD of 0.72Å(Figure 3, which is well below the 2Åthreshold, confirms accurate reproduction of the experimental pose) [55]. This establishes the robustness and predictive reliability of the docking workflow.

**Figure 3:**
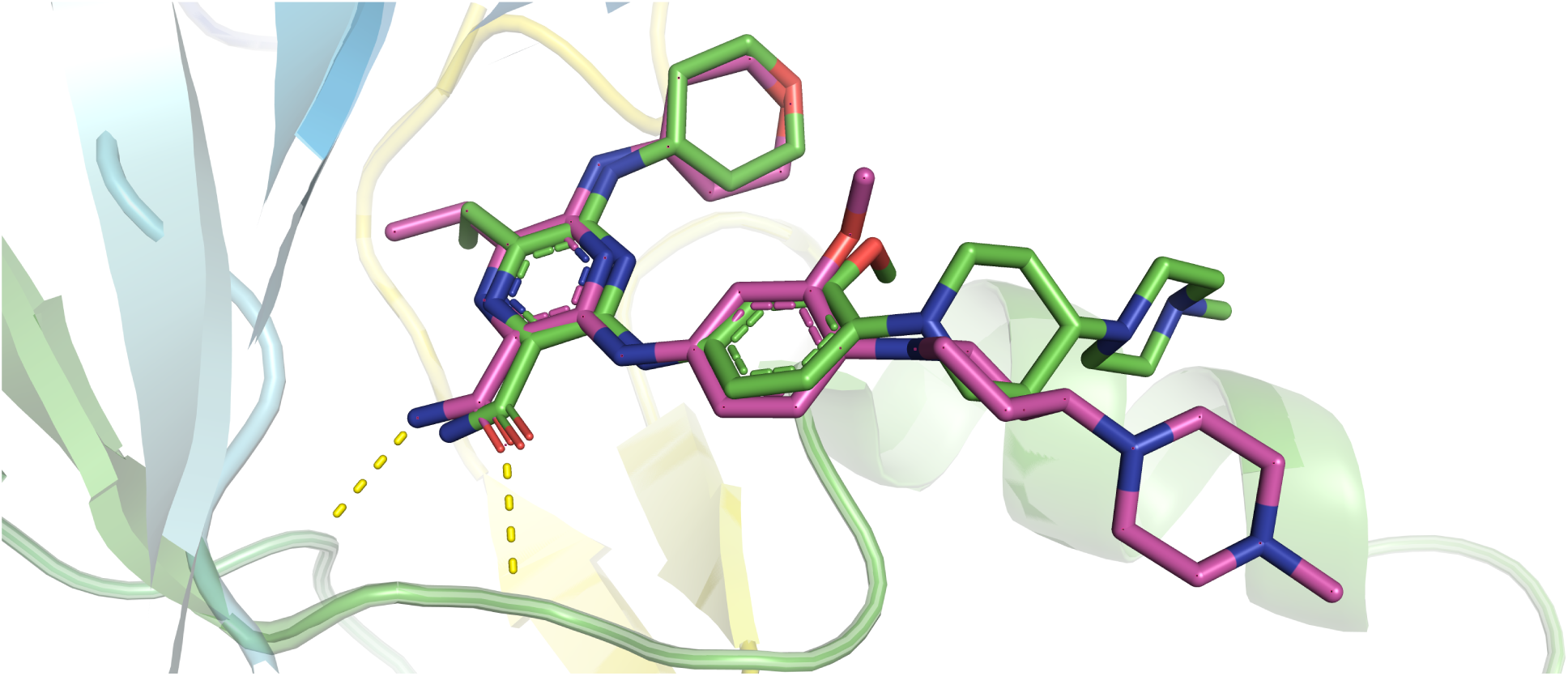
Superposition of native ligand (green) before with the best molecular docking conformations (pink).

### Molecular Interaction Analysis

The top two compounds, MolPort-002-705-878 and MolPort-007-550-904 with binding affinities of –11.33 kcal/mol and –10.66 kcal/mol correspondingly (Table 2), were selected as leads for detailed interaction analysis. MolPort-002-705-878 exhibited multiple key interactions within the active site. It formed four conventional hydrogen bonds with CYS694 (1.69Å), ASP698 (2.15Å), GLU692 (2.05Å), and GLY697 (2.48Å) as shown in Figure 4. A carbon–hydrogen bond was observed with GLY617 (2.59Å), along with a pi–sigma interaction with LEU818 (4.62Å) and an amide–pi stacked interaction with GLY831 (4.03Å). Hydrophobic interactions included pi-alkyl contacts with LEU832 (4.92Å) and GLY831 (3.45Å), and alkyl interactions involving VAL624 (5.11, 5.32Å), VAL675 (4.12Å), ALA642 (4.30Å), LEU818 (5.09, 4.62Å), PHE691 (3.58Å), and LYS644 (4.62Å). These interactions stabilize the ligand within the binding pocket, highlighting its high affinity, with the presence of multiple hydrogen bonds further contributing to its enhanced binding energy [56].

**Figure 4:**
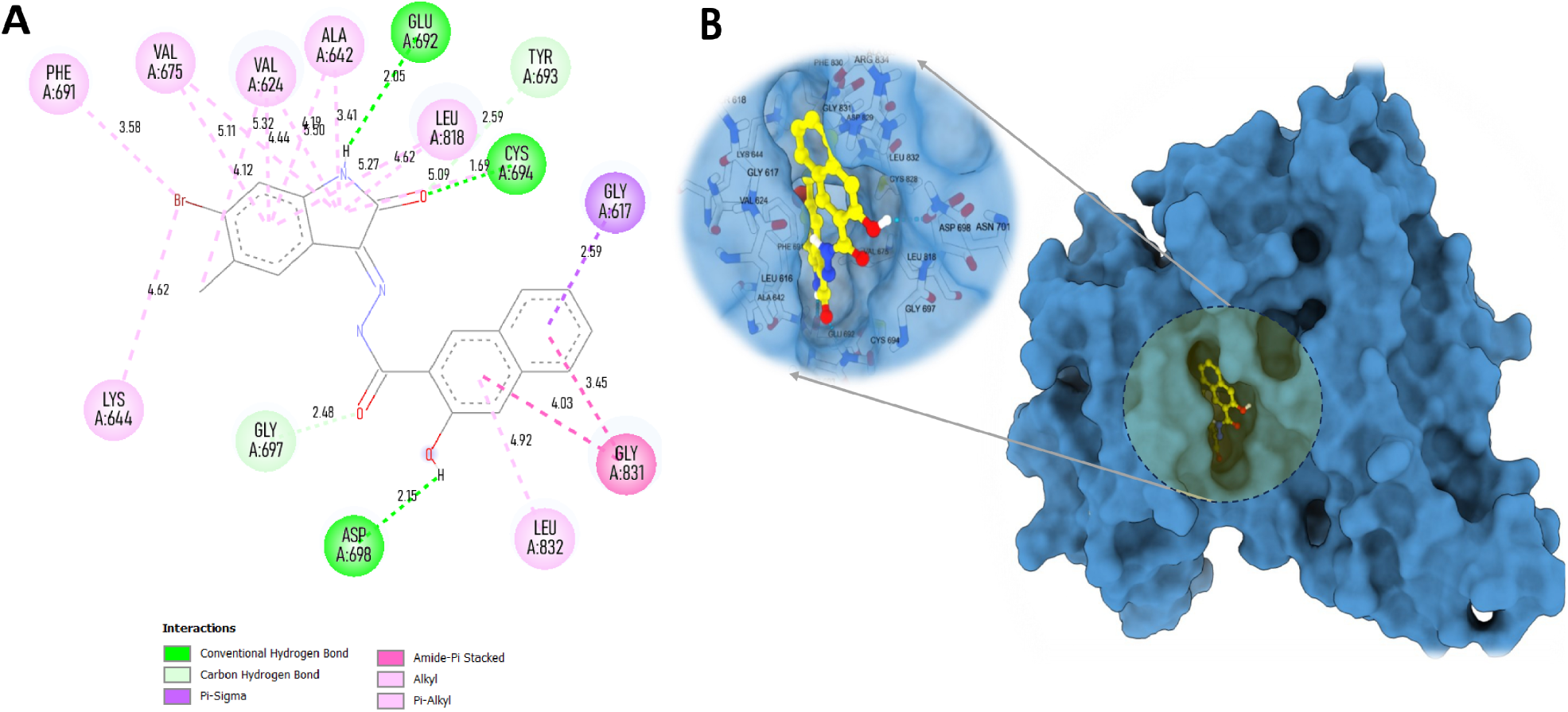
Interaction plots of MolPort-002-705-878: (A) 2D interaction diagram and (B) 3D interaction surface within the active site.

MolPort-007-550-904 also displayed significant interactions within the active sites (Figure 5) forming conventional hydrogen bonds with ASP698 (2.19, 2.21Å), CYS694 (2.77Å), GLU692 (1.77Å), and GLY697 (2.45Å) (Table 3). A carbon-hydrogen bond was formed with LEU832 (2.56Å). Pi-pi T-shaped stacking was observed with PHE691 (4.91Å), and amide-pi stacked interactions was detected with GLY831 (3.69Å) and PHE691 (5.19Å). Pi-alkyl interactions were found with VAL675 (5.20Å), VAL624 (5.03Å), ALA642 (4.46Å), and LEU818 (4.82Å), while alkyl interactions involved LEU616 (3.70Å) and GLY831 (3.69Å). Collectively, these interactions suggest a strong and specific binding orientation in the active site and contend for potent inhibitor of FLT3.

**Figure 5:**
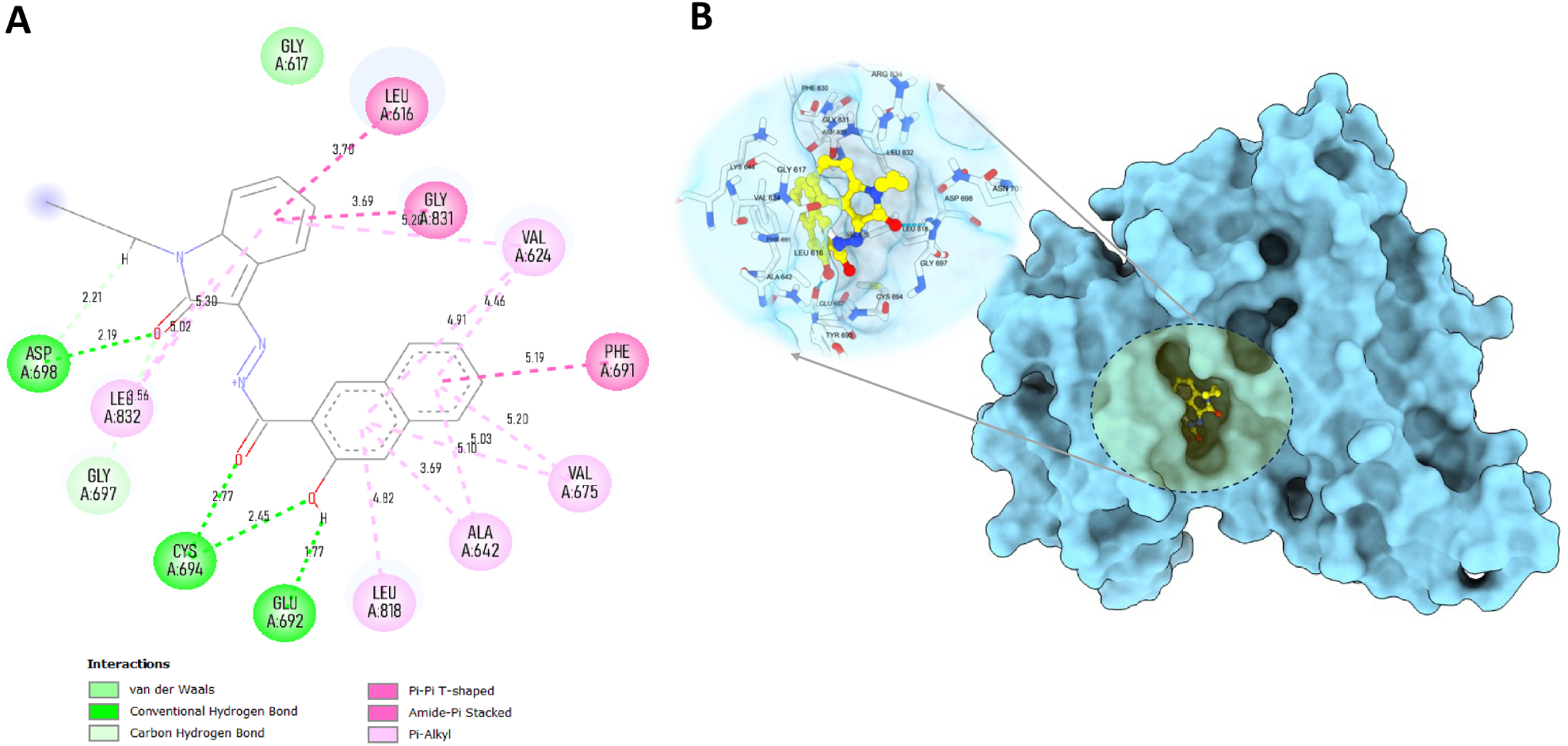
Interaction plots of MolPort-007-550-904: (A) 2D interaction diagram and (B) 3D interaction surface within the active site.

**Table 3:** Interaction analysis of two lead compounds with FLT3 protein.

| Compound | Interaction Types | Interacting Amino Acids (Distance in Å) |
| --- | --- | --- |
| MolPort-002-705-878 | Conventional Hydrogen Bond | CYS694 (1.69), ASP698 (2.15), GLU692 (2.05), GLY697 (2.48) |
|  | Carbon Hydrogen Bond | GLY617 (2.59) |
|  | Pi-Sigma | LEU818 (4.62) |
|  | Amide-Pi Stacked | GLY831 (4.03) |
|  | Pi-Alkyl | LEU832 (4.92), GLY831 (3.45) |
|  | Alkyl | VAL624 (5.11, 5.32), VAL675 (4.12), ALA642 (4.30), LEU818 (5.09, 4.62), PHE691 (3.58), LYS644 (4.62) |
| MolPort-007-550-904 | Conventional Hydrogen Bond | ASP698 (2.19, 2.21), CYS694 (2.77), GLU692 (1.77), GLY697 (2.45) |
|  | Carbon Hydrogen Bond | LEU832 (2.56) |
|  | Pi-Pi T-shaped | PHE691 (4.91) |
|  | Amide-Pi Stacked | GLY831 (3.69), PHE691 (5.19) |
|  | Pi-Alkyl | VAL675 (5.20), VAL624 (5.03), ALA642 (4.46), LEU818 (4.82) |
|  | Alkyl | LEU616 (3.70), GLY831 (3.69) |

### MD Simulations

The RMSD values of the protein fluctuated between 1.2–1.5Åfor the MolPort-007-550-904 complex and approximately 2.5–3.0Åfor the MolPort-002-705-878 complex during molecular dynamics simulations (Figures 6A and 7A), indicating overall structural stability. The ligands, however, exhibited distinct dynamic behaviors: MolPort-007-550-904 showed greater mobility (RMSD ranging from 2.0–3.0Å), suggestive of conformational flexibility, whereas MolPort-002-705-878 exhibited a more restrained profile (1.5–2.0Å), reflecting enhanced binding stability.

**Figure 6:**
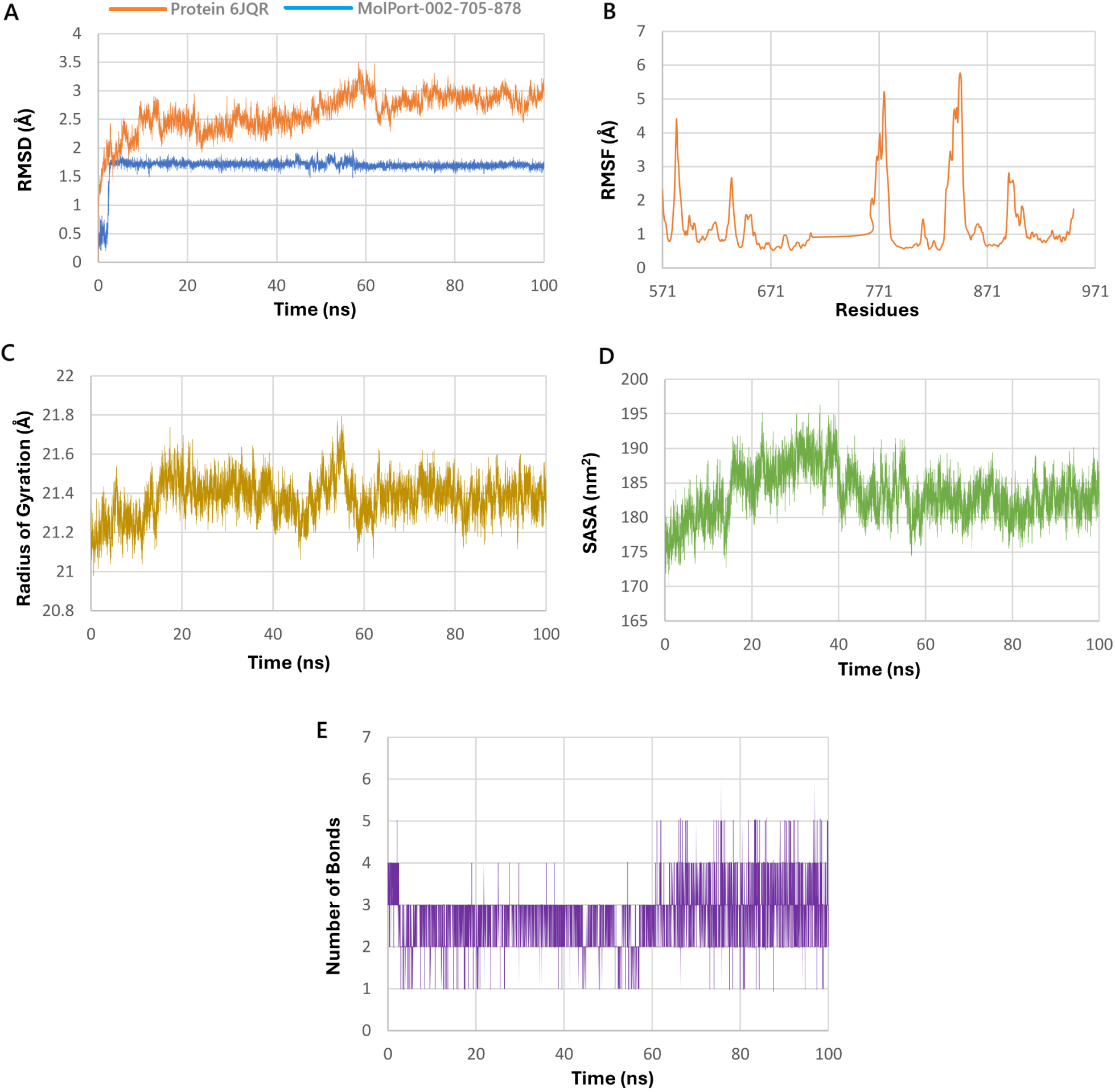
Molecular dynamics simulation analysis of the 6JQR & MolPort-002-705-878 complex over 100 ns. (A) RMSD plot comparing structural stability of the protein (orange) and ligand (blue). (B) RMSF plot showing residue-specific flexibility of the protein, with higher fluctuations in loop regions. (C) Radius of gyration (Rg) indicating consistent compactness of the protein. (D) SASA plot reflecting solvent accessibility changes during the simulation. (E) Hydrogen bonding dynamics over time showing consistent ligand–protein interactions, reaching up to six bonds at peak points. These results suggest strong and stable ligand binding with enhanced interaction stability.

**Figure 7:**
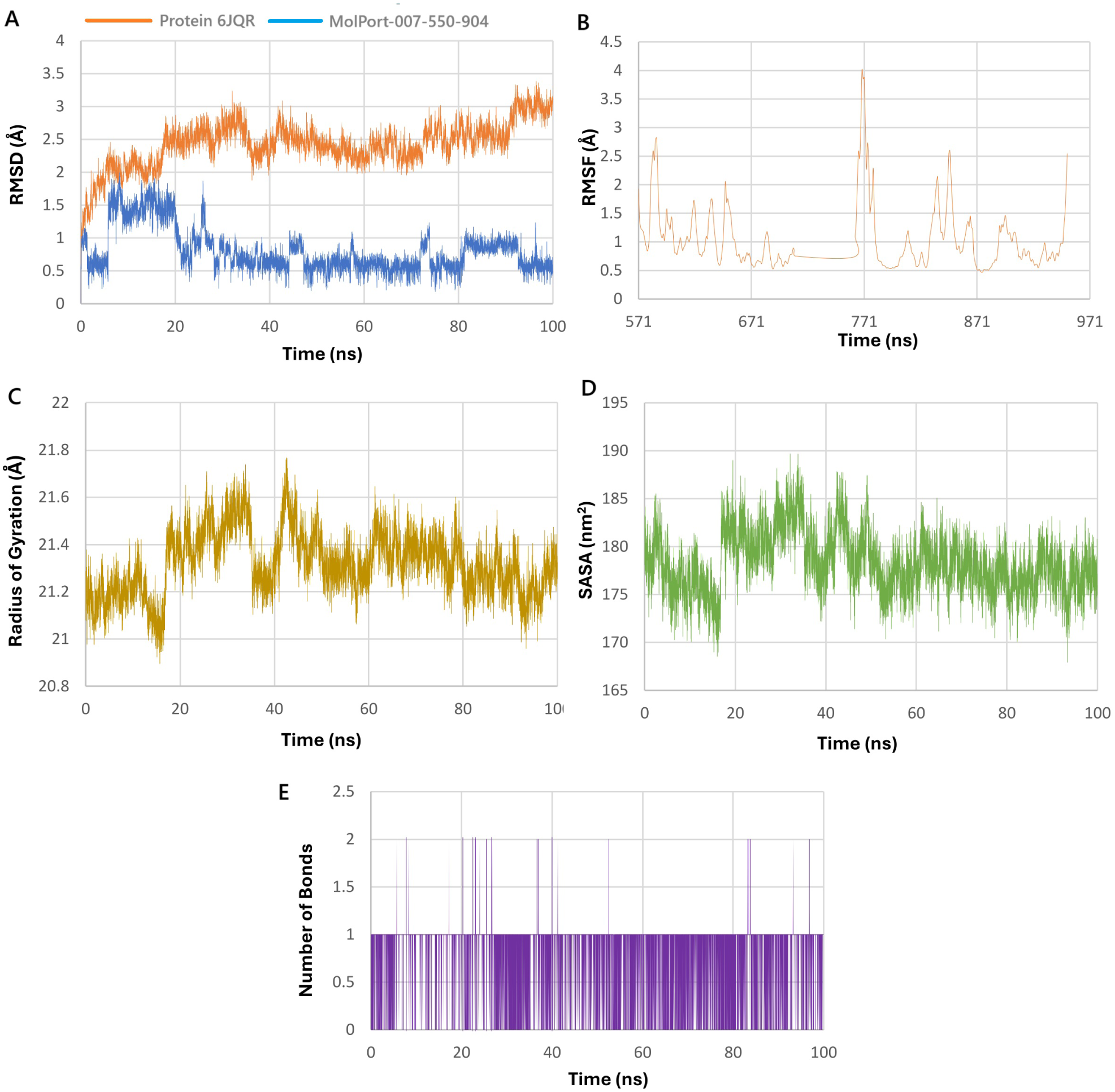
Molecular dynamics simulation analysis of the 6JQR & MolPort-007-550-904 complex over 100 ns. (A) RMSD plot showing backbone stability of the protein (blue) and ligand mobility (orange). (B) RMSF plot representing residue-wise fluctuations of the protein backbone. (C) Radius of gyration (Rg) profile showing the compactness of the protein structure. (D) Solvent-accessible surface area (SASA) illustrating changes in surface exposure. (E) Number of hydrogen bonds formed between the ligand and protein over the simulation time. The data indicate moderate ligand flexibility and intermittent hydrogen bonding, with the protein maintaining overall structural stability.

RMSF (Root Mean Square Fluctuation) analyses (Figures 6B and 7B) showed that while the majority of residues fluctuated below 2.0Å, certain regions in the MolPort-002-705-878 complex exhibited fluctuations reaching up to 6.0Å, indicating localized flexibility. The radius of gyration (Rg) remained consistent in both simulations, with values fluctuating narrowly around 21.0–21.7Å(Figures 6C and 7C), suggesting maintenance of a compact protein fold.

SASA (Solvent Accessible Surface Area) trends (Figures 6D and 7D) indicated a slight decrease for the MolPort-007-550-904 complex, whereas MolPort-002-705-878 showed a modest increase, reflecting minor structural adjustments. Hydrogen bond analysis (Figures 6E and 7E) revealed that the MolPort-007-550-904 complex formed 1–2 transient hydrogen bonds on average, while the MolPort-002-705-878 complex consistently maintained 3–4 bonds, occasionally peaking at 6, implying a more stable and potentially higher-affinity interaction within the ligand-binding site of the protein [20].

MolPort-002-705-878 forms a more stable and persistent complex with 6JQR, as evidenced by lower ligand RMSD, stronger hydrogen bonding, and increased solvent exposure, whereas MolPort-007-550-904 exhibits greater conformational mobility and weaker stabilizing interactions, indicating a more transient binding mode [57].

### MM/GBSA Binding Free Energy

The van der Waals (VDWAALS) and electrostatic (EEL) contributions for MolPort-002-705-878 were –43.83 kcal/mol and –19.74 kcal/mol, respectively, while for MolPort-007-550-904 they were –41.01 kcal/mol and –3.08 kcal/mol. The polar solvation energy (EGB) was 29.00 kcal/mol for MolPort-002-705-878 and 22.24 kcal/mol for MolPort-007-550-904, and non-polar solvation (ESURF) contributions were –5.14 kcal/mol and –5.18 kcal/mol, respectively. Gas-phase interaction (GGAS) and solvation energies (GSOLV) were –63.57/24.34 kcal/mol for MolPort-002-705-878 and –44.08/17.05 kcal/mol for MolPort-007-550-904. Collectively, these components yielded binding free energies (Δ*G*) of –39.23 kcal/mol and –27.03 kcal/mol, confirming stronger overall binding for MolPort-002-705-878. The MM/GBSA energy profiles (Figure 8, Table 4) align with dynamic and structural stability observed in simulations, reinforcing the drug-like potential of both ligands [21].

**Figure 8:**
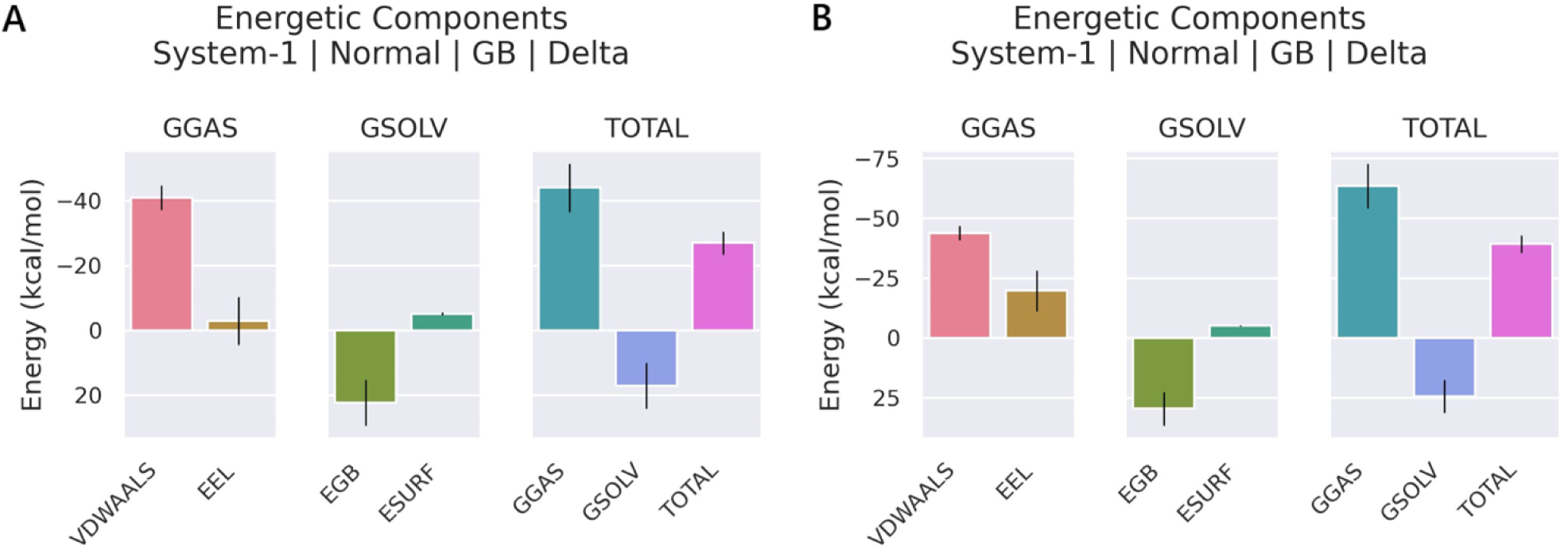
MMGBSA binding energy component analysis; (A) MolPort-007-550-904 and (B) MolPort-002-705-878.

**Table 4:** MMGBSA Binding Energy Components.

| Energy Component | MolPort-002-705-878 | MolPort-007-550-904 |
| --- | --- | --- |
| VDWAALS | -43.83 | -41.01 |
| EEL | -19.74 | -3.08 |
| EGB | 29.48 | 22.24 |
| ESURF | -5.14 | -5.18 |
| GGAS | -63.57 | -44.08 |
| GSOLV | 24.34 | 17.05 |
| TOTAL $\Delta G$ | **-39.23** | **-27.03** |

### Principal Component Analysis (PCA)

#### Eigenvalues of the Covariance Matrix

The eigenvalue spectrum as shown in Panel A of Figure 9 illustrates the distribution of atomic fluctuations along the principal components. Both systems, 6JQR–(MolPort-002-705-878) complex (red) and 6JQR–(MolPort-007-550-904) complex (orange), exhibit a steep decline in the first few eigen-values, followed by a long tail of lower-magnitude components. This trend indicates that the majority of the protein’s collective motions are captured within the first few principal components. The first complex displays slightly higher initial eigenvalues, suggesting stronger collective motions and potentially greater conformational variability compared to that of the latter complex.

**Figure 9:**
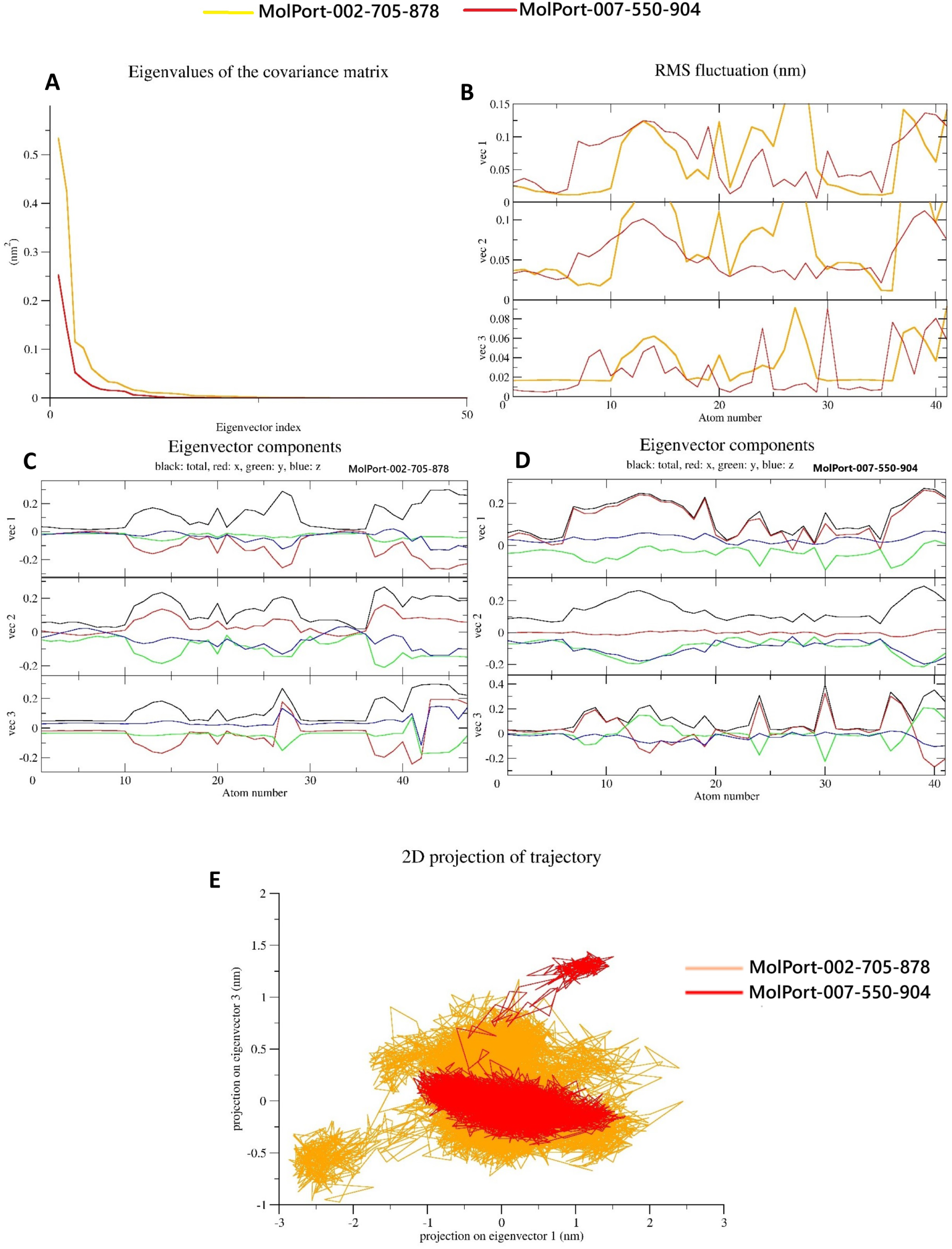
(A) Eigenvalue distribution (B) RMS fluctuation (C,D) Eigenvector Components (E) 2D projection of the trajectory for both the leads.

**Figure 10:**
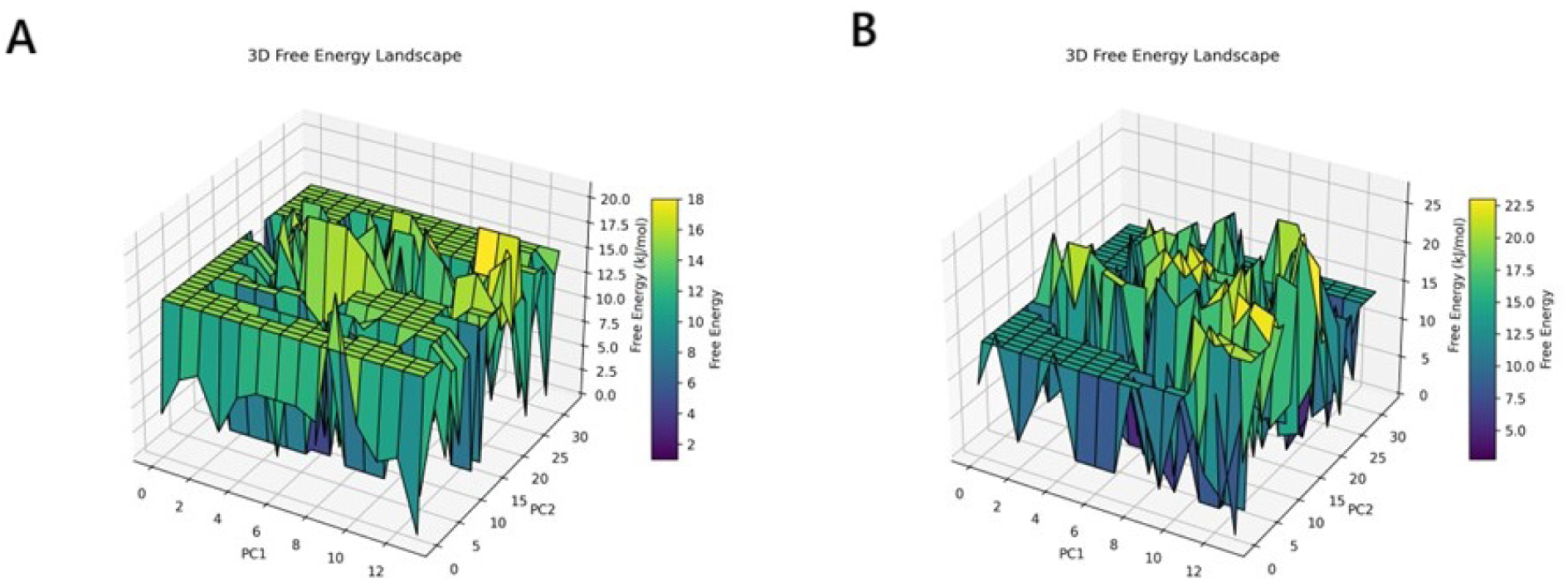
Free Energy Landscape 3D of (A) 6JQR–(MolPort-007-550-904) (B) 6JQR–(MolPort-002-705-878).

#### RMS Fluctuations Along Principal Components

Panel B presents the root mean square (RMS) fluctuation of atomic positions along the first three principal eigenvectors (vec1, vec2, and vec3). The 6JQR–(MolPort-007-550-904) complex exhibits greater amplitude fluctuations, particularly along vec1 and vec3, with peaks exceeding 0.1 nm. These pronounced fluctuations imply more dynamic movements across key residues. In contrast, the 6JQR–(MolPort-002-705-878) complex demonstrates lower atomic fluctuations, indicative of a more rigid structure with constrained deviations along the principal axes [58].

#### Eigenvector Component Analysis

Panels C (for the 6JQR–MolPort-002-705-878 complex) and D (for the 6JQR–MolPort-007-550-904 complex) illustrate the x, y, and z components of the first three eigenvectors. In both systems, the total motion (black) is predominantly governed by the x-component (red), followed by moderate contributions from the y-(green) and z-(blue) directions. However, the MolPort-007-550-904 complex displays greater directional variation, especially in vec1 and vec2, indicating more spatially diverse and extended motion. The MolPort-002-705-878 complex shows more uniform directional behavior, suggesting a compact and directionally consistent dynamic profile [58].

## 2D Projection of Trajectory

Panel E provides a two-dimensional projection of the atomic motions along the first two principal components (vec1 vs. vec2). The 6JQR–(MolPort-007-550-904) complex (in orange) explores a wider and more dispersed conformational space, indicating the sampling of multiple structural states during the simulation. Conversely, the 6JQR–(MolPort-002-705-878) complex (in red) remains tightly clustered within a narrow conformational region, reflecting greater structural rigidity and limited flexibility. This projection emphasizes the dynamic plasticity of the MolPort-007-550-904 complex relative to the more stable and conformationally constrained behavior of the MolPort-002-705-878 complex, which may correlate with differences in functional or binding characteristics [19].

### Free Energy Landscape (FEL)

FEL analysis of the 6JQR–(MolPort-007-550-904) complex (Panel A of Figure 2) exhibits a rugged topology characterized by multiple shallow and deep minima dispersed across the PC1–PC2 conformational space. This distribution indicates frequent transitions among diverse conformational states, suggesting a higher degree of structural flexibility and the presence of several semi-stable intermediates sampled throughout the simulation [59]. In contrast, the FEL of the 6JQR–(MolPort-002-705-878) complex (Panel B of Figure 2) presents a more compact yet energetically heterogeneous landscape. While several peaks and troughs are evident, the overall topology is more condensed, featuring fewer and narrower basins. The presence of deep energy wells suggests that the system adopts fewer, but more stable, conformational states compared to the MolPort-007-550-904 complex.

### Quantum Properties Evaluation

#### Geometry Optimization

The optimized molecular geometries of MolPort-002-705-878 and MolPort-007-550-904 are illustrated in Figure 11 and, key geometric and electronic parameters are summarized in Table 5. MolPort-002-705-878 exhibits a significantly lower global minimum energy (–3734.930 Hartree) than MolPort-007-550-904 (–1240.047 Hartree), indicating greater thermodynamic stability, while both molecules show geometrically stable conformations as supported by their very low RMS Cartesian force values [60]. The computed dipole moments of MolPort-002-705-878 and MolPort-007-550-904 are 8.027 Debye and 6.411 Debye, respectively. These relatively high dipole moments suggest that both compounds are capable of engaging in dipole–dipole interactions with protein residues, which can further influence the binding affinity of the ligand–protein complexes [61].

**Figure 11:**
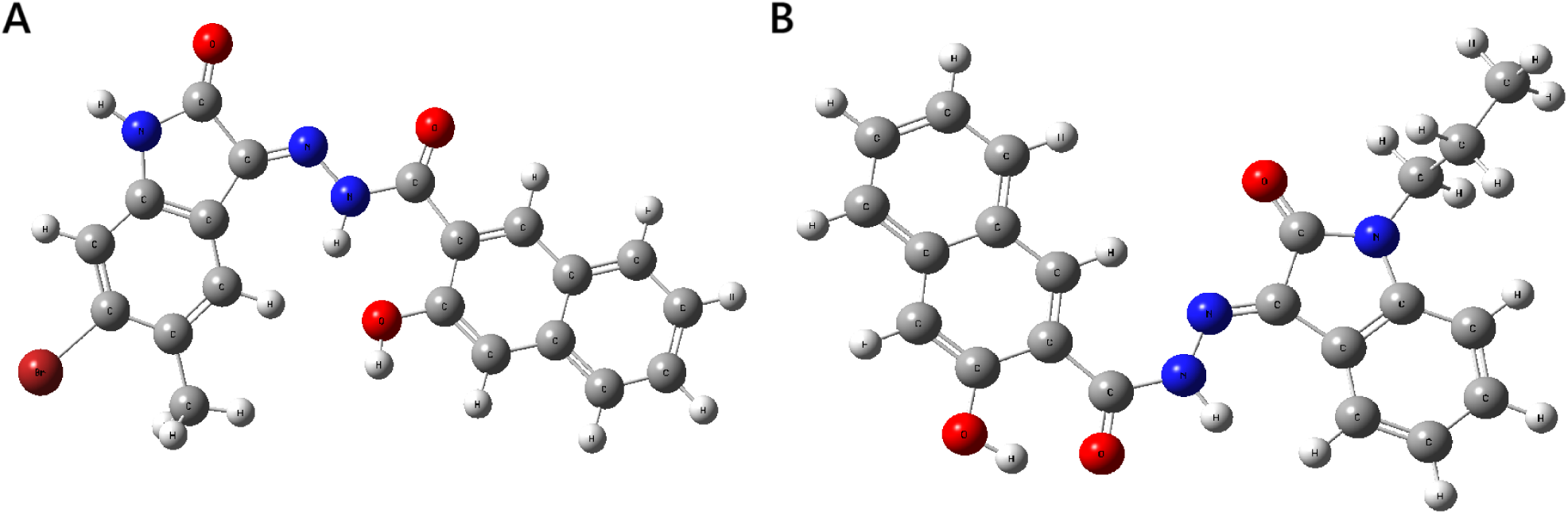
The optimized molecular geometries of the (A) MolPort-002-705-878 and (B) MolPort-007-550-904 molecules.

**Table 5:**
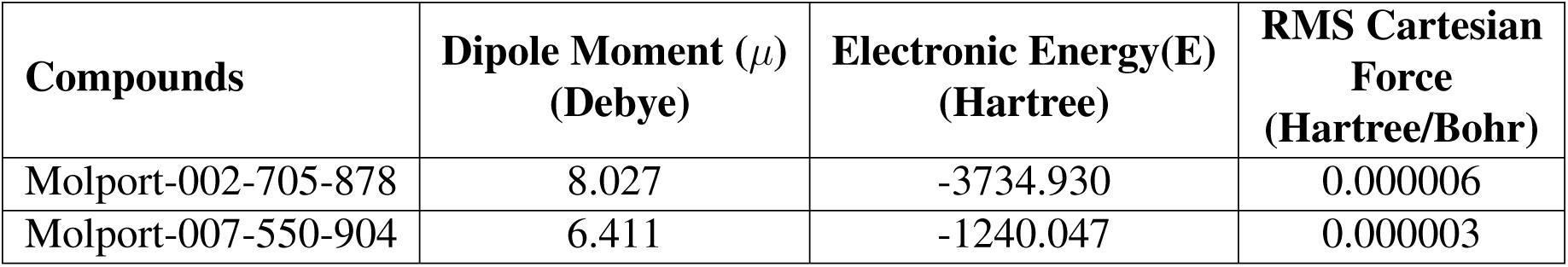
Calculated electronic energy, dipole moment, and RMS Cartesian force of lead compounds.

| Compounds | Dipole Moment ($\mu$)<br>(Debye) | Electronic Energy(E)<br>(Hartree) | RMS Cartesian<br>Force<br>(Hartree/Bohr) |
| --- | --- | --- | --- |
| Molport-002-705-878 | 8.027 | -3734.930 | 0.000006 |
| Molport-007-550-904 | 6.411 | -1240.047 | 0.000003 |

#### Frontier Molecular orbitals (FMOs)

Frontier molecular orbitals (FMOs) offer qualitative understanding of the likelihood of electron transfer between the HOMO (highest occupied molecular orbital) and the LUMO (lowest unoccupied molecular orbital). Among the two leads, MolPort-007-550-904 exhibits the highest HOMO energy (–0.208 eV) and lowest LUMO energy (–0.100 eV), while MolPort-002-705-878 shows the lowest HOMO (–0.227 eV) and highest LUMO (–0.0933 eV), indicating their differing electron-donating and -accepting tendencies. Difference of HUMO and LUMO, also known as energy gaps, for MolPort-007-550-904 and MolPort-002-705-878 are calculated to be 0.108 eV and 0.134 eV, correspondingly. MolPort-007-550-904 has shown a smaller energy gap which implies greater chemical softness and higher reactivity, facilitating more efficient electron transfer compared to MolPort-002-705-878s [62]. The spatial distributions of the HOMO and LUMO are presented in the Figure 12.

**Figure 12:**
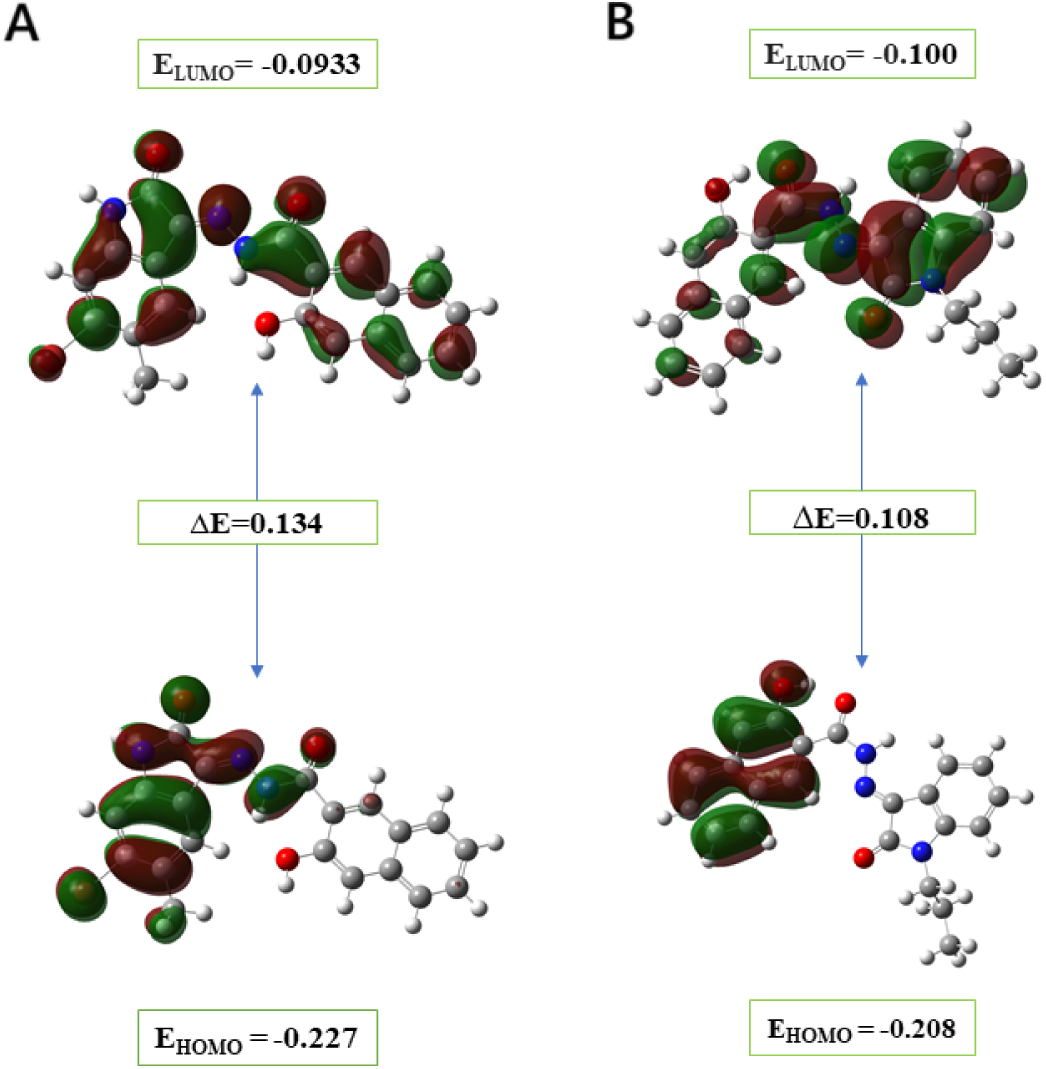
Frontier molecular orbitals (HOMO and LUMO) at the ground state for (A) MolPort-002-705-878 and (B) MolPort-007-550-904.

#### Global Reactivity Parameters

The parameters describing reactivity of compounds are reported in Table 6. Chemical hardness and electrophilicity index values are positive for both molecules, indicating their potential to modulate binding energy in protein–ligand interactions and engage in charge transfer processes [63]. The low chemical potential (*µ*) values for MolPort-007-550-904 (−0.154 eV) and MolPort-002-705-878 (−0.160 eV) further suggest a tendency to donate electrons within the biological environment [64]. MolPort-007-550-904 exhibits greater softness (18.518) compared to MolPort-002-705-878 (14.970), suggesting higher polarizability and reactivity. The higher electronegativity and chemical hardness values of MolPort-002-705-878 indicate relatively stronger electronic stability and lower susceptibility to perturbation than MolPort-007-550-904 [65]. Collectively, these descriptors imply that while both molecules are chemically active, MolPort-007-550-904 may exhibit stronger interaction potential through more facile electron transfer.

**Table 6:** Computed global reactivity descriptors (in eV) for two leads.

| Parameter | MolPort-002-705-878 | MolPort-007-550-904 |
| --- | --- | --- |
| Ionization energy (I) | 0.227 | 0.208 |
| Electron affinity (A) | 0.093 | 0.100 |
| Chemical hardness ($\eta$) | 0.069 | 0.054 |
| Chemical Softness ($\sigma$) | 14.970 | 18.518 |
| Electronegativity ($\chi$) | 0.160 | 0.154 |
| Chemical potential ($\mu$) | -0.160 | -0.154 |
| Electrophilicity ($\omega$) | 0.192 | 0.219 |

#### Molecular Electrostatic Potential (MEP)

The color-mapped surface of MEP facilitates the clear visualization of the electrostatic potential, where gradients indicate the intensity of local charge distribution. The color scheme follows an order where red comes first, blue comes last, and orange, yellow, and green lie in the middle in that sequence. Negative potential regions (red to yellow) typically correspond to electron-rich zones favorable for electrophilic attack, whereas blue regions indicate electron-deficient, electropositive sites that tend to engage with nucleophiles [66]. Green represents areas of near-zero potential. As depicted in Figure 13, regions of negative electrostatic potential are primarily localized around electronegative atoms such as nitrogen and oxygen, indicating potential nucleophilic sites for interactions. In contrast, zones of positive potential are situated near hydrogen atoms, suggesting electrophilic character, while carbon atoms exhibit a mixed pattern depending on their local chemical environment. These surface regions, prone to nucleophilic and electrophilic attacks, offer insight into possible interaction modes with biological macromolecules, as they may complement the electrostatic landscape of the protein binding pocket [67].

**Figure 13:**
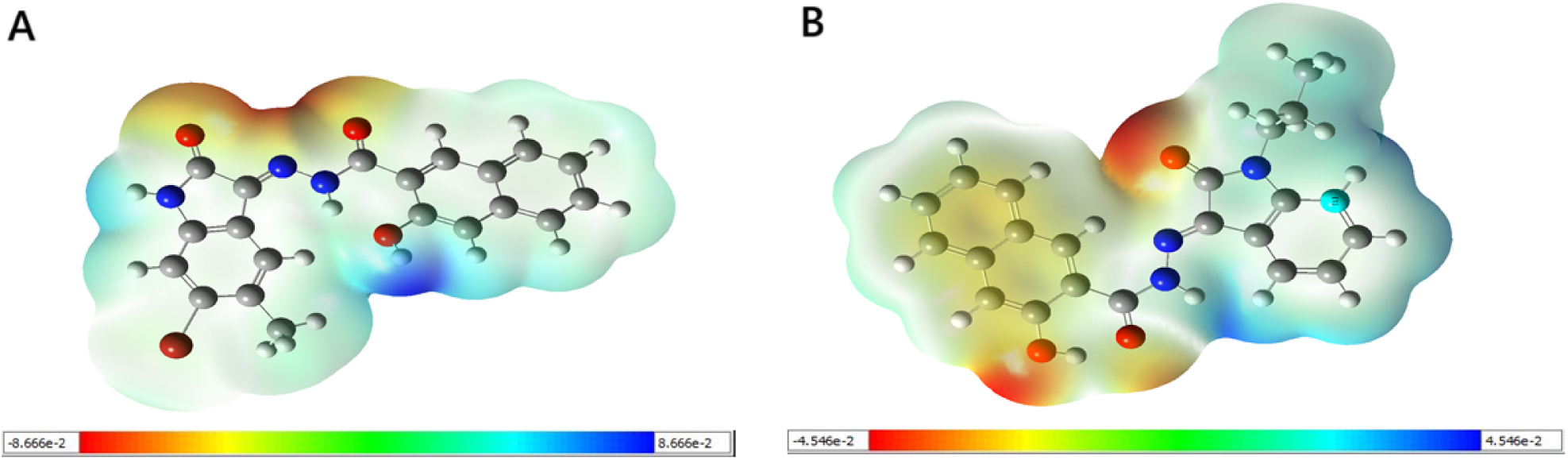
MEP maps with various color coding on the isodensity surface for (A) Molport-002-705-878 and (B) Molport-007-550-904.

#### Natural bond orbital analysis (NBO)

NBO analysis revealed significant intramolecular charge transfer (ICT) interactions in both lead compounds, contributing to their electronic stability (Table 3S & 4S). In MolPort-002-705-878, the dominant donor–acceptor transition was LP(1)N6 *→ π*^∗^(C3–O19), with a stabilization energy of 48.74 kcal/mol. Other notable interactions included LP(1)N13 *→ π*^∗^(C8–O20), LP(1)N6 *→ π*^∗^(C4– C12), and LP(2)O19 *→ σ*^∗^(C3–N6), with energies ranging from 28.74 to 46.92 kcal/mol. In MolPort-007-550-904, a remarkably strong LP(1)C14 *→ π*^∗^(C5–C9) interaction was observed, with an *E*(2) value of 84.40 kcal/mol, alongside additional stabilizing transitions from LP(1)N3 and LP(1)N11 into conjugated *π*^∗^ orbitals. These high second-order perturbation energies (*E*(2)) highlight extensive electron delocalization and resonance stabilization within the lead molecules, particularly through lone pair to *π*^∗^ and *σ*^∗^ interactions [51]. Such hyperconjugative effects suggest increased molecular rigidity and thermodynamic stability, which may translate into enhanced binding affinity and pharmacophoric stability in protein–ligand complexes [67]. The conjugation patterns, especially in heteroatom-rich environments, support the hypothesis that these molecules possess well-distributed electronic densities conducive to hydrogen bonding, *π*–*π* stacking, and electrostatic complementarity with the FLT3 active site [68].

#### Mulliken and Natural Population Analyses

The evaluation of Mulliken and natural population charges is widely performed in quantum chemical studies to identify reactive sites, model electrostatic interactions, and support force field parameterization in molecular simulations [69]. A detailed tabulation of both charge types is presented in Table 2S, and a bar chart comparing them is shown in Figure 14.

**Figure 14:**
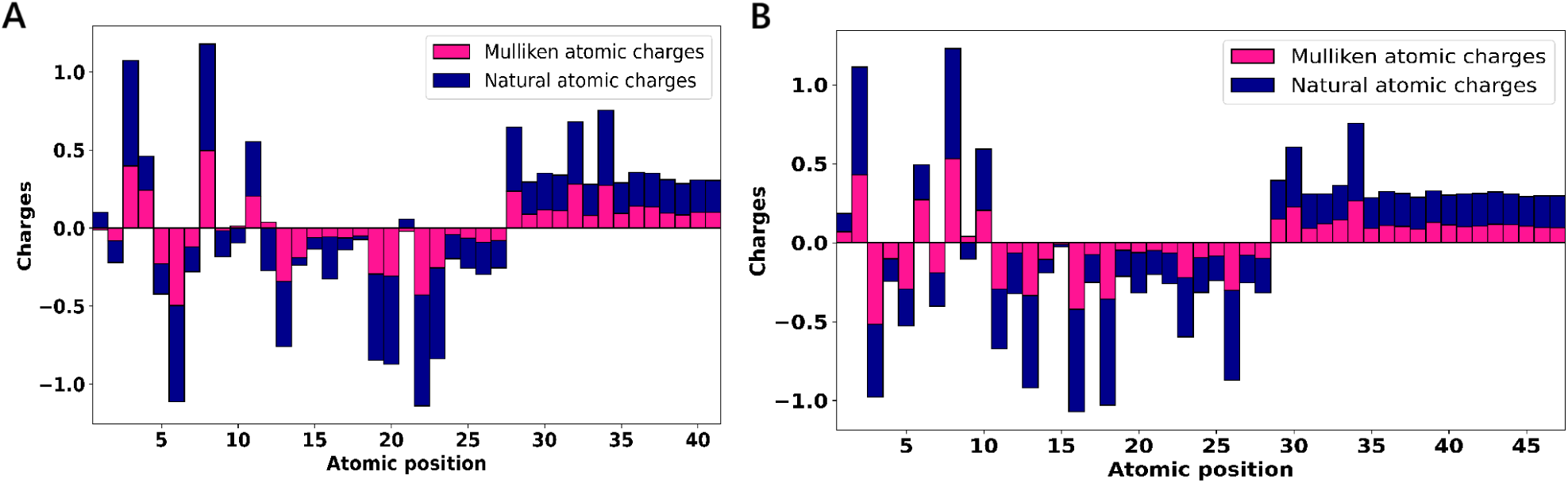
Mulliken and natural atomic charges of (A) MolPort-002-705-878 and (B) MolPort-007-550-904. Charges are stacked when of the same sign; opposite signs are plotted in diverging directions from zero.

According to Mulliken and NBO-based natural charge analyses, the most positive charge is localized on the C8 atom in both lead compounds, indicating a region of electron deficiency (Table 2S). N3 and O18 in MolPort-007-550-904, and N6 and O22 in MolPort-002-705-878, exhibited the most negative Mulliken and natural charges, respectively, reflecting substantial electron localization at key electronegative centers with potential binding relevance [70]. These sites were also involved in interactions during docking analysis, consistent with charge-based predictions.

In MolPort-002-705-878, the bromine atom (Br21) exhibited a small positive Mulliken charge and a positive natural charge, indicating reduced electronic reactivity compared to typical halogen atoms.

This atypical behavior may arise from electron delocalization or shielding by adjacent functional groups [71]. Among hydrogen atoms, the highest Mulliken charge values were observed on H34 in MolPort-007-550-904 and H32 in MolPort-002-705-878, suggesting potential involvement in electrostatic interactions or hydrogen bonding [72]. Carbon atoms showed a distribution of both positive and negative charges, reflecting their diverse electronic environments.

### ADMET Analysis

The pharmacokinetic and toxicological characteristics of the lead compounds were evaluated through comprehensive *in silico* analyses to assess their potential as drug-like molecules suitable for preclinical development. MolPort-002-705-878 exhibits a favorable absorption profile, with high predicted gastrointestinal (GI) permeability and compliance with key criteria for drug-likeness, as set by Lipinski, Ghose, Veber, and Egan rules. The compound has a moderate lipophilicity profile (consensus LogP = 3.47), topological polar surface area (TPSA) of 90.79 Å ^2^ and three rotatable bonds, which supports passive diffusion across biological membranes [25, 73]. Despite being moderately soluble by ESOL and poorly soluble by AliLogS and Silicos-IT models, its drug-likeness is reinforced by favorable bioavailability (score: 0.55) and minimal synthetic complexity. Importantly, the compound is predicted to be non-substrate for P-glycoprotein and lacks blood–brain barrier permeability (Figure 2S). The predicted Lethal dose 50 (LD_50_) for this lead is approximately 3009 mg/kg, corresponding to toxicity class V under the Globally Harmonized System (GHS), indicative of low acute toxicity [74]. It is estimated to be non-carcinogenic, non-mutagenic, and non-cytotoxic. Organ-specific toxicity predictions indicate potential hepatotoxicity, nephrotoxicity, and neurotoxicity, while cardiotoxicity and respiratory toxicity are unlikely. Additionally, molecular initiating event (MIE) analyses suggest activation of the aryl hydrocarbon receptor (AhR) and potential mitochondrial membrane disruption, both of which may warrant further investigation during safety profiling.

MolPort-007-550-904, similarly, demonstrates high predicted GI absorption and complies with all major drug-likeness rules, aside from a single Muegge violation. It shows slightly higher structural flexibility, with five rotatable bonds and a TPSA of 82.00 Å ^2^. Although its consensus LogP (3.46) and solubility predictions mirror the first compound, it’s slightly elevated fraction of sp^3^ carbons may suggest better metabolic stability [75]. As with MolPort-002-705-878, it is not BBB-permeant and avoids P-glycoprotein-mediated efflux (Figure 2S). However, it exhibits a broader CYP inhibition profile *in silico*, inhibiting CYP1A2, CYP2C19, CYP2C9, and CYP3A4, raising possible concerns regarding metabolic interactions. In toxicity evaluation, this lead has comparatively lower LD_50_ of 1400 mg/kg and is classified as toxicity class IV, suggesting moderate acute toxicity [74]. It exhibits a low probability of causing carcinogenicity, immunotoxicity, or endocrine disruption. However, it shows higher toxicity liabilities overall, with predicted neurotoxicity, nephrotoxicity, respiratory toxicity, and mutagenicity. While activation of nuclear receptor pathways appears minimal, this compound also engages in mitochondrial toxicity mechanisms and should be evaluated carefully by *in vitro* assays.

## 4 Conclusion

This investigation focuses on FLT3, aiming to discover innovative small molecules with significant therapeutic potential. In this study, the structure-based screening tool Pharmit and the comprehensive commercial chemical library MolPort were leveraged to maximize chemical diversity and increase the likelihood of identifying highly potent and selective FLT3 inhibitors. The rigorous process began with an initial virtual screening using Pharmit, followed by docking-based screening of 419 selected hits, resulting in the identification of two promising lead molecules: MolPort-002-705-878 and MolPort-007-550-904. The additional validation of these leads were performed using molecular dynamics simulations to probe stability of protein-ligand binding, density functional theory (DFT) exploration to characterize quantum chemical behavior, and ADMET profiling to evaluate drug-likeness, all of which collectively support these lead as potential FLT3 antagonists. However, certain aspects identified in the ADMET analysis necessitate experimental validation and further clinical investigation. This strategy exemplifies a scalable, data-driven framework for next-generation drug discovery that can be adapted to investigate diverse biomolecular targets across a broad range of diseases. While the approach shows significant promise, it also presents limitations, such as high computational demands, reliance on in silico predictions without experimental confirmation, and the potential incompleteness of commercial chemical databases. To overcome these challenges, future efforts should integrate high-performance computing infrastructure, foster interdisciplinary collaboration with experimental pharmacology laboratories for empirical validation, and incorporate emerging non-commercial compound repositories. As the field evolves, harmonizing computational insights with experimental rigor will be essential for transforming virtual hits into viable clinical candidates.

## 5 Supplementary Information

This paper reports the key outcomes of our investigation. Further data and additional images are provided in the Supplementary Information.

