## Supplemental file for "Structure-Guided Discovery and Characterization of Novel FLT3 Inhibitors for Acute Myeloid Leukemia Treatment"

### Supplementary Information:

Table 1S: MolPort Compound IDs selected for docking after pharmacophore filtering.

| Compounds | Compounds | Compounds | Compounds |
|---|---|---|---|
| MolPort-002-705-878 | MolPort-046-857-239 | MolPort-001-028-598 | MolPort-002-655-342 |
| MolPort-007-903-602 | MolPort-002-329-300 | MolPort-000-732-736 | MolPort-019-911-361 |
| MolPort-019-910-904 | MolPort-003-714-345 | MolPort-002-945-344 | MolPort-006-844-396 |
| MolPort-007-550-904 | MolPort-001-529-307 | MolPort-002-664-858 | MolPort-019-707-345 |
| MolPort-007-606-024 | MolPort-001-544-997 | MolPort-019-911-366 | MolPort-002-037-067 |
| MolPort-004-856-891 | MolPort-001-539-548 | MolPort-008-288-055 | MolPort-002-629-529 |
| MolPort-001-650-645 | MolPort-004-856-888 | MolPort-008-286-984 | MolPort-002-664-450 |
| MolPort-001-671-202 | MolPort-006-804-052 | MolPort-002-486-321 | MolPort-004-850-504 |
| MolPort-000-817-743 | MolPort-001-521-829 | MolPort-001-495-409 | MolPort-002-666-266 |
| MolPort-004-856-885 | MolPort-000-225-262 | MolPort-001-539-207 | MolPort-002-668-549 |
| MolPort-004-846-897 | MolPort-004-942-287 | MolPort-047-920-301 | MolPort-002-563-446 |
| MolPort-047-019-496 | MolPort-001-610-192 | MolPort-046-703-376 | MolPort-018-580-482 |
| MolPort-001-647-167 | MolPort-046-689-686 | MolPort-002-672-426 | MolPort-002-667-910 |
| Molport-051-903-815 | MolPort-002-668-584 | MolPort-000-431-547 | MolPort-001-645-325 |
| MolPort-051-700-067 | MolPort-002-665-607 | MolPort-002-667-942 | MolPort-002-668-107 |
| MolPort-002-664-137 | MolPort-002-671-622 | MolPort-001-644-819 | MolPort-051-755-753 |
| MolPort-002-938-003 | MolPort-002-046-193 | MolPort-001-549-987 | Molport-051-838-512 |
| MolPort-000-851-496 | MolPort-000-858-348 | MolPort-002-989-879 | MolPort-002-665-252 |
| MolPort-001-620-913 | MolPort-002-664-182 | MolPort-002-251-242 | MolPort-002-134-370 |
| MolPort-007-987-586 | MolPort-002-667-049 | MolPort-044-435-979 | MolPort-002-667-454 |
| MolPort-002-566-143 | MolPort-004-878-609 | MolPort-001-981-118 | MolPort-002-817-646 |
| MolPort-001-639-043 | MolPort-001-639-878 | MolPort-002-212-894 | MolPort-000-884-203 |
| MolPort-007-662-228 | MolPort-002-139-081 | MolPort-001-954-874 | MolPort-002-134-368 |
| MolPort-000-225-285 | MolPort-018-542-219 | MolPort-035-946-294 | MolPort-003-143-942 |
| MolPort-004-850-680 | MolPort-002-838-441 | MolPort-051-700-066 | MolPort-002-212-495 |
| MolPort-047-355-235 | MolPort-002-139-019 | MolPort-009-652-536 | MolPort-002-539-711 |
| MolPort-002-507-745 | MolPort-002-563-305 | MolPort-002-665-369 | MolPort-002-666-245 |
| MolPort-002-663-032 | MolPort-002-668-772 | MolPort-002-564-370 | Molport-051-892-177 |
| MolPort-002-247-103 | MolPort-002-607-129 | MolPort-002-563-677 | MolPort-002-153-986 |
| MolPort-002-735-434 | MolPort-039-050-507 | MolPort-002-251-118 | MolPort-001-912-640 |
| MolPort-002-469-922 | MolPort-002-666-462 | MolPort-002-666-949 | MolPort-002-134-369 |
| MolPort-049-225-129 | MolPort-002-668-414 | MolPort-004-850-679 | MolPort-038-402-714 |
| MolPort-000-859-344 | MolPort-001-649-761 | MolPort-001-539-825 | MolPort-023-332-859 |
| MolPort-019-913-459 | MolPort-003-846-724 | MolPort-002-507-746 | MolPort-000-765-456 |
| MolPort-000-555-518 | MolPort-002-470-753 | MolPort-002-668-265 | MolPort-004-933-308 |

| Compounds | Compounds | Compounds | Compounds |
|---|---|---|---|
| MolPort-002-507-747 | MolPort-002-547-197 | MolPort-002-599-502 | MolPort-001-544-405 |
| MolPort-000-650-811 | MolPort-002-468-571 | MolPort-001-586-477 | MolPort-002-566-100 |
| MolPort-002-602-189 | MolPort-001-893-495 | MolPort-002-667-070 | MolPort-002-668-973 |
| MolPort-002-665-240 | MolPort-002-139-031 | MolPort-002-902-191 | MolPort-001-490-415 |
| MolPort-001-620-497 | MolPort-047-156-478 | MolPort-002-640-643 | MolPort-045-967-963 |
| MolPort-002-566-538 | MolPort-002-003-681 | MolPort-002-247-093 | MolPort-002-635-812 |
| MolPort-000-934-045 | MolPort-003-802-422 | MolPort-002-565-489 | MolPort-047-933-048 |
| MolPort-019-797-234 | MolPort-002-585-457 | MolPort-002-602-969 | MolPort-002-667-991 |
| MolPort-002-745-815 | MolPort-002-601-356 | MolPort-002-664-939 | MolPort-002-567-705 |
| MolPort-002-604-045 | MolPort-002-673-970 | MolPort-002-816-077 | MolPort-002-603-395 |
| MolPort-000-921-623 | MolPort-003-355-489 | MolPort-004-788-067 | MolPort-002-345-146 |
| MolPort-047-586-059 | MolPort-027-352-440 | MolPort-002-667-688 | Molport-004-802-782 |
| Molport-051-901-940 | MolPort-002-587-437 | MolPort-002-507-743 | MolPort-049-220-517 |
| MolPort-001-014-891 | MolPort-002-003-758 | MolPort-002-600-113 | MolPort-002-565-702 |
| MolPort-009-753-114 | MolPort-001-610-110 | MolPort-002-568-090 | MolPort-000-919-403 |
| MolPort-002-605-666 | MolPort-047-798-457 | MolPort-003-875-129 | MolPort-002-136-586 |
| MolPort-035-689-865 | MolPort-002-600-911 | MolPort-000-851-136 | MolPort-004-262-677 |
| MolPort-002-048-223 | MolPort-000-224-828 | Molport-051-810-669 | MolPort-004-942-293 |
| MolPort-002-602-566 | MolPort-002-250-837 | MolPort-002-663-314 | MolPort-001-955-424 |
| MolPort-002-121-860 | MolPort-035-763-685 | MolPort-002-705-687 | MolPort-046-510-951 |
| MolPort-002-604-343 | MolPort-048-722-691 | MolPort-004-345-557 | MolPort-000-801-443 |
| MolPort-004-350-349 | MolPort-002-798-643 | Molport-004-802-783 | MolPort-002-563-493 |
| MolPort-002-568-543 | MolPort-002-173-815 | MolPort-046-791-775 | MolPort-028-914-300 |
| MolPort-002-635-848 | MolPort-035-685-038 | MolPort-010-466-524 | MolPort-002-251-549 |
| MolPort-000-678-736 | MolPort-002-625-922 | MolPort-015-143-278 | MolPort-022-375-119 |
| MolPort-000-734-890 | MolPort-028-957-496 | MolPort-051-765-205 | MolPort-035-689-869 |
| Molport-051-804-083 | MolPort-029-940-689 | MolPort-002-666-483 | MolPort-000-482-961 |
| MolPort-028-750-277 | MolPort-000-149-495 | MolPort-003-818-711 | MolPort-039-322-891 |
| MolPort-007-899-726 | MolPort-004-330-336 | MolPort-047-970-305 | MolPort-046-528-459 |
| MolPort-016-625-795 | MolPort-047-970-407 | MolPort-000-145-535 | MolPort-002-816-887 |
| MolPort-000-521-655 | MolPort-002-748-021 | MolPort-046-934-151 | MolPort-009-196-316 |
| MolPort-002-603-739 | MolPort-001-661-776 | MolPort-044-553-661 | MolPort-002-563-576 |
| MolPort-023-330-468 | MolPort-014-880-695 | MolPort-042-646-687 | MolPort-049-242-540 |
| MolPort-002-858-741 | MolPort-044-832-228 | MolPort-000-931-718 | MolPort-046-194-590 |
| Molport-051-804-543 | Molport-045-019-282 | MolPort-002-566-888 | MolPort-047-967-835 |
| MolPort-047-967-884 | MolPort-047-740-535 | MolPort-001-772-772 | MolPort-002-319-269 |
| MolPort-035-785-556 | MolPort-002-462-096 | MolPort-035-874-381 | MolPort-047-969-126 |
| MolPort-035-394-359 | MolPort-000-146-043 | MolPort-004-363-487 | MolPort-013-209-953 |
| MolPort-004-293-855 | MolPort-000-885-524 | MolPort-002-461-971 | MolPort-039-332-433 |

| Compounds | Compounds | Compounds | Compounds |
|---|---|---|---|
| MolPort-000-271-683 | MolPort-001-899-637 | MolPort-047-970-273 | MolPort-027-679-828 |
| MolPort-000-149-357 | MolPort-002-471-423 | MolPort-000-163-937 | MolPort-051-766-167 |
| MolPort-002-889-632 | MolPort-046-940-595 | MolPort-002-563-218 | MolPort-019-930-873 |
| MolPort-046-522-317 | MolPort-003-920-584 | MolPort-002-041-254 | MolPort-004-955-729 |
| MolPort-038-438-241 | MolPort-006-716-553 | MolPort-022-379-265 | MolPort-001-790-840 |
| MolPort-002-462-058 | MolPort-002-462-059 | MolPort-002-628-504 | MolPort-027-948-434 |
| MolPort-001-761-481 | Molport-051-912-158 | MolPort-000-147-051 | MolPort-046-934-207 |
| MolPort-001-771-862 | MolPort-002-566-786 | MolPort-001-649-791 | MolPort-022-898-151 |
| MolPort-046-860-003 | MolPort-049-202-162 | MolPort-002-568-945 | MolPort-051-649-859 |
| MolPort-015-141-656 | MolPort-039-203-876 | MolPort-038-434-884 | MolPort-003-904-445 |
| MolPort-004-363-488 | MolPort-047-413-669 | MolPort-051-544-861 | MolPort-000-704-473 |
| MolPort-006-716-291 | MolPort-044-183-544 | MolPort-004-291-541 | MolPort-023-330-328 |
| MolPort-002-041-266 | MolPort-000-271-686 | MolPort-023-330-096 | MolPort-044-817-891 |
| MolPort-044-558-710 | MolPort-003-749-417 | MolPort-000-271-687 | MolPort-042-681-224 |
| MolPort-047-970-797 | MolPort-039-203-878 | MolPort-004-767-950 | MolPort-019-878-674 |
| MolPort-009-196-794 | MolPort-016-578-683 | MolPort-038-386-980 | MolPort-038-521-591 |
| MolPort-027-948-360 | MolPort-009-199-618 | MolPort-003-929-057 | MolPort-046-686-088 |
| MolPort-035-690-033 | MolPort-001-814-813 | MolPort-005-311-290 | MolPort-003-847-744 |
| MolPort-051-589-588 | MolPort-042-672-242 | MolPort-009-199-188 | MolPort-051-767-274 |
| MolPort-020-248-041 | MolPort-047-413-648 | MolPort-000-210-149 | MolPort-011-016-227 |
| MolPort-047-413-628 | MolPort-023-332-377 | MolPort-016-582-338 | MolPort-016-579-028 |
| MolPort-004-767-393 | MolPort-006-710-144 | MolPort-027-947-944 | MolPort-035-687-287 |
| MolPort-004-768-362 | MolPort-004-949-903 | MolPort-023-222-588 | MolPort-023-331-591 |
| MolPort-004-812-210 | MolPort-044-557-618 | MolPort-002-719-266 | MolPort-046-196-178 |
| Molport-051-891-697 | MolPort-019-879-186 | MolPort-020-222-334 | MolPort-000-210-144 |
| MolPort-000-146-587 | MolPort-047-352-420 | MolPort-004-816-169 | MolPort-020-217-041 |
| MolPort-008-000-002 | MolPort-046-194-946 | MolPort-004-768-381 | MolPort-001-893-025 |
| MolPort-016-583-176 | MolPort-019-996-089 | MolPort-004-767-492 | MolPort-000-884-453 |
| MolPort-008-646-580 | MolPort-020-003-025 | MolPort-047-598-283 | MolPort-002-121-685 |
| MolPort-000-720-481 | MolPort-046-424-804 | MolPort-000-841-538 | MolPort-022-628-246 |
| MolPort-039-137-277 | MolPort-004-942-582 | MolPort-004-891-628 | |

Table 2S: Mulliken and Natural charges of all the atoms in MolPort-007-550-904 and MolPort-002-705-878 molecules.

| **Atoms of MolPort-007-550-904** | **Mulliken charges for MolPort-007-550-904** | **Natural charges for MolPort-007-550-904** | **Atoms of MolPort-002-705-878** | **Mulliken charges for MolPort-002-705-878** | **Natural charges for MolPort-002-705-878** |
|---|---|---|---|---|---|
| C1 | 0.068716 | 0.11626 | C1 | -0.012779 | 0.09953 |
| C2 | 0.428234 | 0.68362 | C2 | -0.08277 | -0.13782 |
| N3 | -0.517527 | -0.45981 | C3 | 0.39672 | 0.67633 |
| C4 | -0.101593 | -0.14344 | C4 | 0.24141 | 0.21741 |
| C5 | -0.293929 | -0.23351 | C5 | -0.230521 | -0.19226 |
| C6 | 0.271755 | 0.22075 | N6 | -0.495991 | -0.61729 |
| N7 | -0.192721 | -0.20910 | N7 | -0.120937 | -0.15896 |
| C8 | 0.530469 | 0.69732 | C8 | 0.497059 | 0.68211 |
| C9 | 0.038490 | -0.10620 | C9 | -0.017922 | -0.16530 |
| C10 | 0.203963 | 0.38884 | C10 | 0.011402 | -0.09650 |
| N11 | -0.292988 | -0.37843 | C11 | 0.202887 | 0.35137 |
| C12 | -0.065163 | -0.25799 | C12 | 0.035656 | -0.27304 |
| O13 | -0.335211 | -0.58370 | N13 | -0.342662 | -0.41759 |
| C14 | -0.106358 | -0.08648 | C14 | -0.191111 | -0.04624 |
| C15 | -0.010996 | -0.01612 | C15 | -0.062872 | -0.07397 |
| O16 | -0.422602 | -0.64951 | C16 | -0.057371 | -0.27046 |
| C17 | -0.076863 | -0.17646 | C17 | -0.064478 | -0.07582 |
| O18 | -0.357808 | -0.67174 | C18 | -0.050274 | -0.02374 |
| C19 | -0.048919 | -0.16487 | O19 | -0.295504 | -0.55213 |
| C20 | -0.063085 | -0.25510 | O20 | -0.308311 | -0.56534 |
| C21 | -0.049889 | -0.15247 | Br21 | -0.021068 | 0.05726 |
| C22 | -0.066952 | -0.19355 | O22 | -0.429384 | -0.71175 |
| C23 | -0.222670 | -0.37462 | C23 | -0.256448 | -0.58249 |
| C24 | -0.094463 | -0.22195 | C24 | -0.041814 | -0.15570 |
| C25 | -0.085509 | -0.15608 | C25 | -0.065941 | -0.19071 |
| C26 | -0.301953 | -0.56848 | C26 | -0.092787 | -0.20424 |
| C27 | -0.079999 | -0.17384 | C27 | -0.080135 | -0.17570 |
| C28 | -0.099169 | -0.21779 | H28 | 0.234754 | 0.41022 |
| H29 | 0.149235 | 0.24413 | H29 | 0.088805 | 0.20499 |
| H30 | 0.225580 | 0.37810 | H30 | 0.116296 | 0.23437 |
| H31 | 0.090826 | 0.21600 | H31 | 0.111699 | 0.22621 |
| H32 | 0.120540 | 0.18825 | H32 | 0.281323 | 0.39805 |
| H33 | 0.142658 | 0.21836 | H33 | 0.079944 | 0.20054 |
| H34 | 0.264133 | 0.49063 | H34 | 0.272516 | 0.48153 |

| Atoms of MolPort-007-550-904 | Mulliken charges for MolPort-007-550-904 | Natural charges for MolPort-007-550-904 | Atoms of MolPort-002-705-878 | Mulliken charges for MolPort-002-705-878 | Natural charges for MolPort-002-705-878 |
|---|---|---|---|---|---|
| H35 | 0.090561 | 0.19354 | H35 | 0.093385 | 0.19665 |
| H36 | 0.108095 | 0.21365 | H36 | 0.140534 | 0.21732 |
| H37 | 0.101297 | 0.20979 | H37 | 0.135991 | 0.21422 |
| H38 | 0.084295 | 0.20208 | H38 | 0.095726 | 0.21422 |
| H39 | 0.128045 | 0.19931 | H39 | 0.084119 | 0.20148 |
| H40 | 0.112272 | 0.18856 | H40 | 0.100388 | 0.20586 |
| H41 | 0.100247 | 0.20849 | H41 | 0.100463 | 0.20414 |
| H42 | 0.104091 | 0.20605 | - | - | - |
| H43 | 0.117383 | 0.20268 | - | - | - |
| H44 | 0.113498 | 0.19365 | - | - | - |
| H45 | 0.103192 | 0.18808 | - | - | - |
| H46 | 0.095293 | 0.20058 | - | - | - |
| H47 | 0.093499 | 0.20255 | - | - | - |

Table 3S: Second-order perturbation analysis of the interaction between donor and acceptor orbitals of compound Molport-007-550-904 in NBO basis.

| Donor NBO(i) | Type | Acceptor NBO(j) | Type | E(2)kcal/mol | E(j)-E(i)a.u. | F(i,j)a.u. |
|---|---|---|---|---|---|---|
| C1-C2 | σ | C2-O13 | σ* | 1.77 | 1.25 | 0.42 |
| | | N3-C17 | | 4.83 | 0.98 | 0.61 |
| | | C4-C19 | | 4.72 | 1.20 | 0.67 |
| | | N7-N11 | | 6.67 | 1.04 | 0.75 |
| C1-C4 | σ | C1-N7 | σ* | 4.10 | 1.26 | 0.064 |
| | | C4-C19 | | 3.96 | 1.24 | 0.063 |
| | | C6-C20 | | 4.18 | 1.25 | 0.065 |
| C1-N7 | σ | C1-C2 | σ* | 1.55 | 1.31 | 0.041 |
| | | C1-C4 | | 3.82 | 1.38 | 0.065 |
| | | C8-N11 | | 3.25 | 1.32 | 0.059 |
| | π | C1-N7 | π* | 2.63 | 0.35 | 0.028 |
| | | C2-O13 | | 12.08 | 0.36 | 0.062 |
| | | C4-C9 | | 9.10 | 0.37 | 0.055 |
| C2 - N3 | σ | C1-N7 | σ* | 1.42 | 1.38 | 0.039 |
| | | N3 -C6 | | 2.05 | 1.22 | 0.045 |
| | | C6 -C20 | | 3.87 | 1.36 | 0.065 |
| C2-O13 | σ | C1-C2 | σ* | 2.07 | 1.46 | 0.050 |
| C2-O13 | π | C1-N7 | π* | 4.17 | 0.37 | 0.038 |
| N3-C6 | σ | C2-N3 | σ* | 1.27 | 1.22 | 0.036 |
| | | C2-O13 | | 3.24 | 1.41 | 0.061 |
| | | C4-C19 | | 2.38 | 1.36 | 0.051 |
| | | C6-C20 | | 2.36 | 1.37 | 0.051 |
| N3-C17 | σ | N3-N6 | σ | 2.06 | 1.18 | 0.044 |
| | | C4-C6 | | 1.74 | 1.27 | 0.042 |
| C4-C6 | σ | C1-C4 | σ* | 2.24 | 1.15 | 0.045 |
| | | C1-N7 | | 5.51 | 1.27 | 0.075 |
| | | N3-C17 | | 4.69 | 1.03 | 0.062 |
| | | C4-C19 | | 4.39 | 1.25 | 0.066 |
| | | C6-C20 | | 4.52 | 1.25 | 0.068 |
| | | C19-H35 | | 2.84 | 1.11 | 0.051 |
| C4-C19 | σ | C1-C4 | σ* | 4.40 | 1.18 | 0.065 |
| | | N3-C6 | | 2.19 | 1.15 | 0.045 |
| | | C4-C6 | | 5.24 | 1.24 | 0.072 |
| | | C19-C24 | | 3.32 | 1.28 | 0.058 |
| | π* | C1-N7 | π* | 19.91 | 0.27 | 0.066 |
| | | C6-C20 | | 21.21 | 0.29 | 0.070 |
| | | C24-C25 | | 17.10 | 0.29 | 0.063 |
| C5-C8 | σ | C5-C9 π* | σ* | 3.56 | 1.27 | 0.060 |

| | | | | | | |
|---|---|---|---|---|---|---|
| C5-C9 | σ | C5-C10 | σ* | 4.23 | 1.19 | 0.064 |
| | | C9-C14 | | 3.56 | 1.28 | 0.060 |
| | | C10-O18 | | 3.26 | 1.09 | 0.053 |
| | | C14-C21 | | 3.01 | 1.25 | 0.055 |
| | π | C8-O10 | π* | 30.83 | 0.24 | 0.078 |
| | | C10-C12 | | 19.92 | 0.30 | 0.069 |
| C5-C10 | σ | C5-C9 π* | σ* | 4.12 | 1.25 | 0.064 |
| | | C8-N11 | | 3.33 | 1.06 | 0.054 |
| | | C10-C12 | | 3.65 | 1.26 | 0.061 |
| C6-C20 | σ | C4-C6 | σ* | 5.06 | 1.24 | 0.071 |
| | | C20-C25 | | 3.36 | 1.28 | 0.059 |
| | π | C4-C19 | π* | 16.26 | 0.29 | 0.062 |
| | | C24-C25 | | 22.59 | 0.29 | 0.073 |
| C9-C14 | σ | C5-C8 | σ* | 3.75 | 1.11 | 0.058 |
| | | C14-C15 | | 4.03 | 1.23 | 0.063 |
| C9-H29 | σ | C5-C10 | σ* | 4.95 | 1.00 | 0.063 |
| | | C14-C15 | | 5.05 | 1.05 | 0.065 |
| C10-C12 | σ | C5-C10 | σ* | 3.92 | 1.19 | 0.061 |
| | | C12-C15 | | 3.70 | 1.28 | 0.061 |
| | π | C5-C9 | π* | 13.58 | 0.28 | 0.056 |
| N11-H30 | σ | C5-C8 | σ* | 3.61 | 1.13 | 0.058 |
| C12-C15 | σ | C14-C15 | σ* | 3.77 | 1.22 | 0.061 |
| C12-H31 | σ | C5-C10 | σ* | 5.01 | 0.99 | 0.063 |
| | | C14-C15 | | 4.79 | 1.05 | 0.063 |
| C14-C15 | σ | C9-C14 | σ* | 3.69 | 1.23 | 0.060 |
| | | C14-C21 | | 3.60 | 1.21 | 0.059 |
| C14-C21 | σ | C9-C14 | σ* | 3.86 | 1.24 | 0.062 |
| | | C14-C15 | | 3.74 | 1.21 | 0.060 |
| C15-C22 | σ | C12-C15 | σ* | 3.60 | 1.24 | 0.060 |
| C17-H33 | σ | N3-C6 | σ* | 5.05 | 0.94 | 0.062 |
| O18-H34 | σ | C10-C12 | σ* | 4.89 | 1.32 | 0.072 |
| C19-H24 | σ | C1-C4 | σ* | 5.19 | 1.17 | 0.070 |
| C19-H35 | σ | C4-C6 | σ* | 4.54 | 1.06 | 0.062 |
| C20-C25 | σ | N3-N6 | σ* | 5.96 | 1.13 | 0.074 |
| C20-H36 | σ | C4-C6 | σ* | 4.84 | 1.05 | 0.064 |
| C21-C28 | σ | C14-C21 | σ* | 3.34 | 1.25 | 0.058 |
| | π | C22-C27 | π* | 18.65 | 0.30 | 0.066 |
| C21-H37 | σ | C14-C15 | σ* | 4.72 | 1.05 | 0.063 |
| C22-C27 | σ | C15-C22 | σ* | 3.46 | 1.26 | 0.059 |
| | π | C21-C28 | π* | 15.61 | 0.30 | 0.061 |
| C22-H38 | σ | C14-C15 | σ* | 4.56 | 1.05 | 0.062 |
| C24-C25 | π | C4-C19 | π* | 22.61 | 0.28 | 0.072 |
| | | C6-C20 | | 16.79 | 0.28 | 0.062 |
| C24-H41 | σ | C4-C19 | σ* | 4.24 | 1.09 | 0.061 |
| C25-H42 | σ | C6-C20 | σ* | 4.06 | 1.10 | 0.060 |

| | | | | | | |
|---|---|---|---|---|---|---|
| N3 | LP (1) | C2-O13 | π* | 54.12 | 0.28 | 0.112 |
| | | C6-C20 | | 42.81 | 0.28 | 0.098 |
| N7 | LP (1) | C1-C4 | σ* | 12.80 | 0.85 | 0.094 |
| N11 | LP (1) | C1-N7 | π* | 37.70 | 0.29 | 0.096 |
| | | C8-O16 | | 51.82 | 0.28 | 0.108 |
| O13 | LP (2) | C1-C2 | σ* | 21.99 | 0.64 | 0.108 |
| | | C2-N3 | | 27.07 | 0.68 | 0.123 |
| C14 | LP (1) | C5-C9 | π* | 84.40 | 0.13 | 0.112 |
| | | C21-C28 | | 56.67 | 0.14 | 0.101 |
| O16 | LP (2) | C5-C8 | σ* | 11.28 | 0.74 | 0.083 |
| | | C8-N11 | | 21.92 | 0.70 | 0.112 |
| | | O18-H34 | | 19.29 | 0.73 | 0.108 |
| O18 | LP (1) | C5-C10 | σ* | 8.04 | 1.06 | 0.082 |
| | LP (2) | C10-C12 | π* | 35.86 | 0.34 | 0.102 |

Table 4S: Second-order perturbation analysis of the interaction between donor and acceptor orbitals of compound MOLPORT-002-705-878 in NBO basis.

| Donor NBO(i) | Type | Acceptor NBO(j) | Type | E(2)kcal/mol | E(j)-E(i)a.u. | F(i,j)a.u. |
|---|---|---|---|---|---|---|
| C1-C2 | σ | C1-N7 | σ* | 3.57 | 1.26 | 0.060 |
| | | C2-C9 | | 4.20 | 1.24 | 0.064 |
| C1-C3 | σ | C2-C9 | σ* | 4.74 | 1.20 | 0.068 |
| | | N7-N13 | | 6.09 | 1.03 | 0.071 |
| C1-N7 | σ | C1-C2 | σ* | 3.69 | 1.37 | 0.064 |
| | π | C2-C9 | π* | 9.78 | 0.35 | 0.056 |
| | | C3-O19 | | 12.46 | 0.36 | 0.062 |
| C2-C4 | σ | C1-N7 | σ* | 5.09 | 1.28 | 0.072 |
| | | C2-C9 | | 4.32 | 1.26 | 0.066 |
| | | C4-C12 | | 4.56 | 1.25 | 0.068 |
| C2-C9 | σ | C1-C2 | σ* | 4.44 | 1.18 | 0.065 |
| | | C2-C4 | | 5.05 | 1.24 | 0.071 |
| | π | C1-N7 | π* | 17.91 | 0.28 | 0.064 |
| | | C4-C12 | | 21.44 | 0.28 | 0.071 |
| | | C14-C15 | | 18.82 | 0.28 | 0.066 |
| C3-N6 | σ | C4-C12 | σ* | 4.17 | 1.37 | 0.068 |
| C3-O19 | π | C1-N7 | π* | 4.17 | 0.38 | 0.038 |
| C4-C12 | σ | C2-C4 | σ* | 5.01 | 1.25 | 0.071 |
| | | C12-C14 | | 4.38 | 1.27 | 0.067 |
| | | C14-Br21 | | 4.86 | 0.81 | 0.056 |
| | π | C2-C9 | π* | 16.06 | 0.30 | 0.062 |
| | | C14-C15 | | 21.86 | 0.29 | 0.072 |
| C5-C10 | σ | C5-C11 | σ* | 4.11 | 1.21 | 0.063 |
| | | C11-O22 | | 3.93 | 1.01 | 0.056 |
| | π | C8-O20 | π* | 19.72 | 0.29 | 0.068 |
| | | C11-C16 | | 22.25 | 0.28 | 0.071 |
| | | C17-C18 | | 14.31 | 0.29 | 0.061 |

| | | | | | | |
|---|---|---|---|---|---|---|
| C5-C11 | σ | C5-C10 | σ* | 4.32 | 1.29 | 0.067 |
| | | C11-C16 | | 4.57 | 1.28 | 0.068 |
| C9-C15 | σ | C1-C2 | σ* | 4.61 | 1.16 | 0.065 |
| | | C2-C9 | | 4.19 | 1.27 | 0.065 |
| | | C14-C15 | | 5.01 | 1.25 | 0.071 |
| | | C14-Br21 | | 5.89 | 0.78 | 0.061 |
| C9-H29 | σ | C2-C4 | σ* | 4.43 | 1.05 | 0.061 |
| | | C14-C15 | | 4.46 | 1.09 | 0.062 |
| C10-H30 | σ | C5-C11 | σ* | 5.17 | 1.02 | 0.065 |
| | | C17-C18 | | 4.84 | 1.05 | 0.064 |
| C11-C16 | σ | C5-C11 | σ* | 5.34 | 1.24 | 0.073 |
| | | C16-C18 | | 4.04 | 1.28 | 0.064 |
| | π | C5-C10 | π* | 13.35 | 0.31 | 0.058 |
| | | C17-C18 | | 17.27 | 0.31 | 0.070 |
| C12-C14 | σ | C4-N6 | σ* | 5.21 | 1.16 | 0.069 |
| | | C4-C12 | | 4.15 | 1.30 | 0.066 |
| C12-H31 | σ | C2-C4 | σ* | 4.52 | 1.06 | 0.062 |
| | | C14-C15 | | 4.63 | 1.09 | 0.064 |
| C14-C15 | σ | C9-C15 | σ* | 4.09 | 1.29 | 0.065 |
| | π | C2-C9 | π* | 19.22 | 0.30 | 0.069 |
| | | C4-C12 | | 17.28 | 0.29 | 0.064 |
| C17-C18 | π | C5-C10 | π* | 21.02 | 0.27 | 0.071 |
| | | C11-C16 | | 16.21 | 0.26 | 0.060 |
| | | C24-C26 | | 16.87 | 0.28 | 0.065 |
| | | C25-C27 | | 15.20 | 0.28 | 0.062 |
| O22-H34 | σ | C5-C11 | σ* | 4.15 | 1.28 | 0.066 |
| | | C14-C15 | | 4.88 | 1.07 | 0.065 |
| C24-C26 | π | C17-C18 | π* | 15.73 | 0.29 | 0.064 |
| | | C25-C27 | | 18.67 | 0.29 | 0.066 |
| C24-H38 | σ | C17-C18 | σ* | 4.74 | 1.05 | 0.063 |
| | | C26-C27 | | 4.74 | 1.05 | 0.063 |
| C25-C27 | σ | C18-C25 | σ* | 3.54 | 1.26 | 0.060 |
| | π | C17-C18 | π* | 17.66 | 0.29 | 0.068 |
| | | C24-C26 | | 16.25 | 0.30 | 0.063 |
| C25-H39 | σ | C17-C18 | σ* | 4.61 | 1.06 | 0.063 |
| C26-H40 | σ | C17-C24 | σ* | 4.49 | 1.06 | 0.062 |
| C27-H41 | σ | C18-C25 | σ* | 4.58 | 1.06 | N6.062 |
| N6 | LP (1) | C3-O19 | π* | 48.74 | 0.29 | 0.109 |
| | | C4-C12 | | 41.50 | 0.29 | 0.099 |
| N7 | LP (1) | C1-C2 | **σ*** | 13.48 | 0.83 | 0.095 |
| | | N13-H32 | | 8.15 | 0.79 | 0.072 |
| N13 | LP (1) | C1-N7 | **π*** | 33.79 | 0.29 | 0.092 |
| | | C8-O20 | | 46.92 | 0.30 | 0.109 |
| O19 | LP (2) | C1-C3 | **σ*** | 22.68 | 0.63 | 0.108 |
| | | C3-N6 | | 28.74 | 0.66 | 0.125 |
| O20 | LP (2) | C5-C8 | σ* | 20.30 | 0.64 | 0.104 |
| | | C8-N13 | | 28.51 | 0.66 | 0.124 |
| | LP (2) | C12-C14 | σ* | 3.04 | 0.85 | 0.045 |

| | | | | | | |
|---|---|---|---|---|---|---|
| Br21 | | C14-C15 | | 3.63 | 0.86 | 0.050 |
| | LP (3) | C14-C15 | π* | 9.74 | 0.31 | 0.054 |
| O22 | LP (1) | C11-C16 | σ* | 5.08 | 1.21 | 0.070 |
| | | N13-H32 | | 7.27 | 1.06 | 0.078 |
| | LP (2) | C11-C16 | π* | 25.00 | 0.38 | 0.091 |

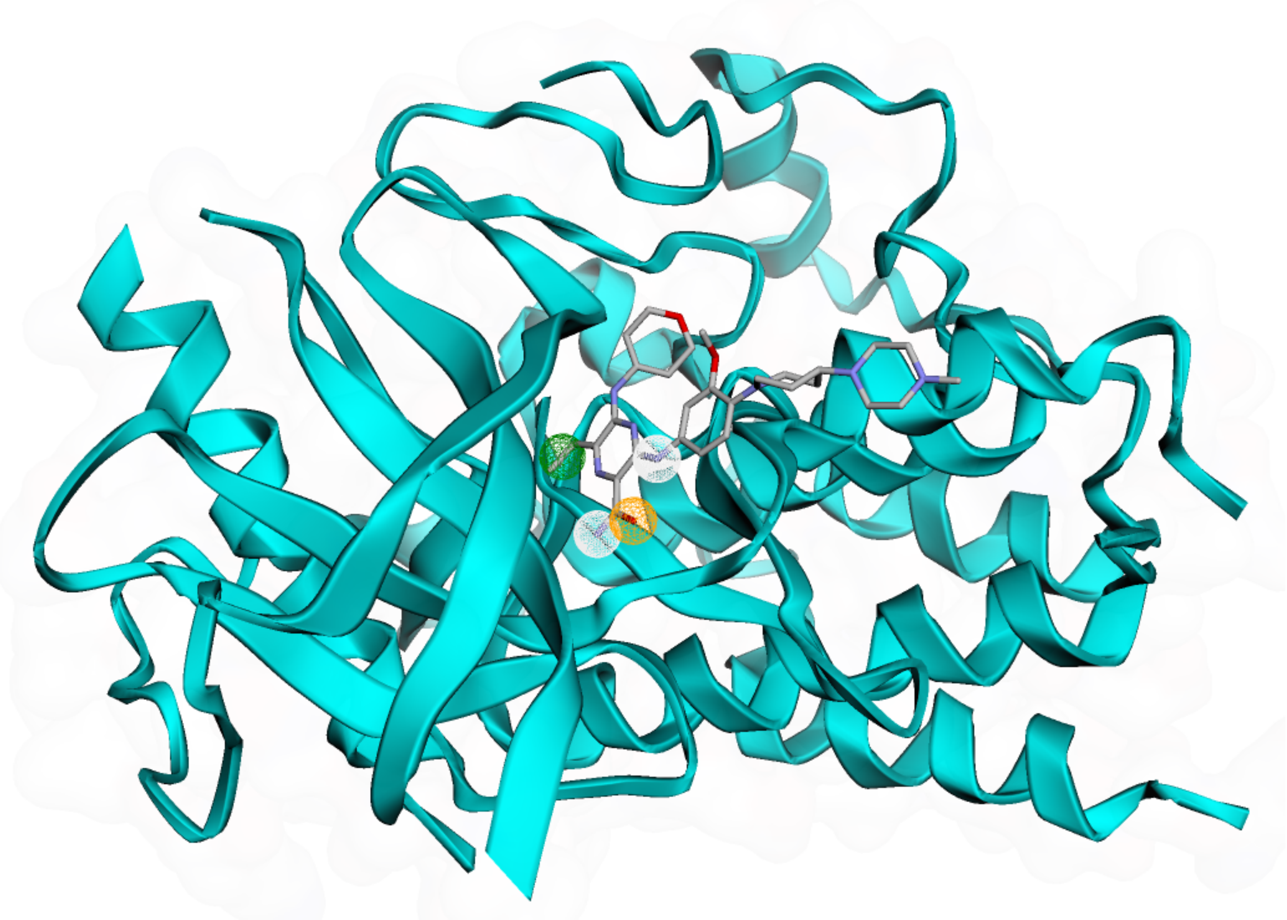

Figure 1S: Generated Pharmacophores in Gliternitib within FLT3 protein.

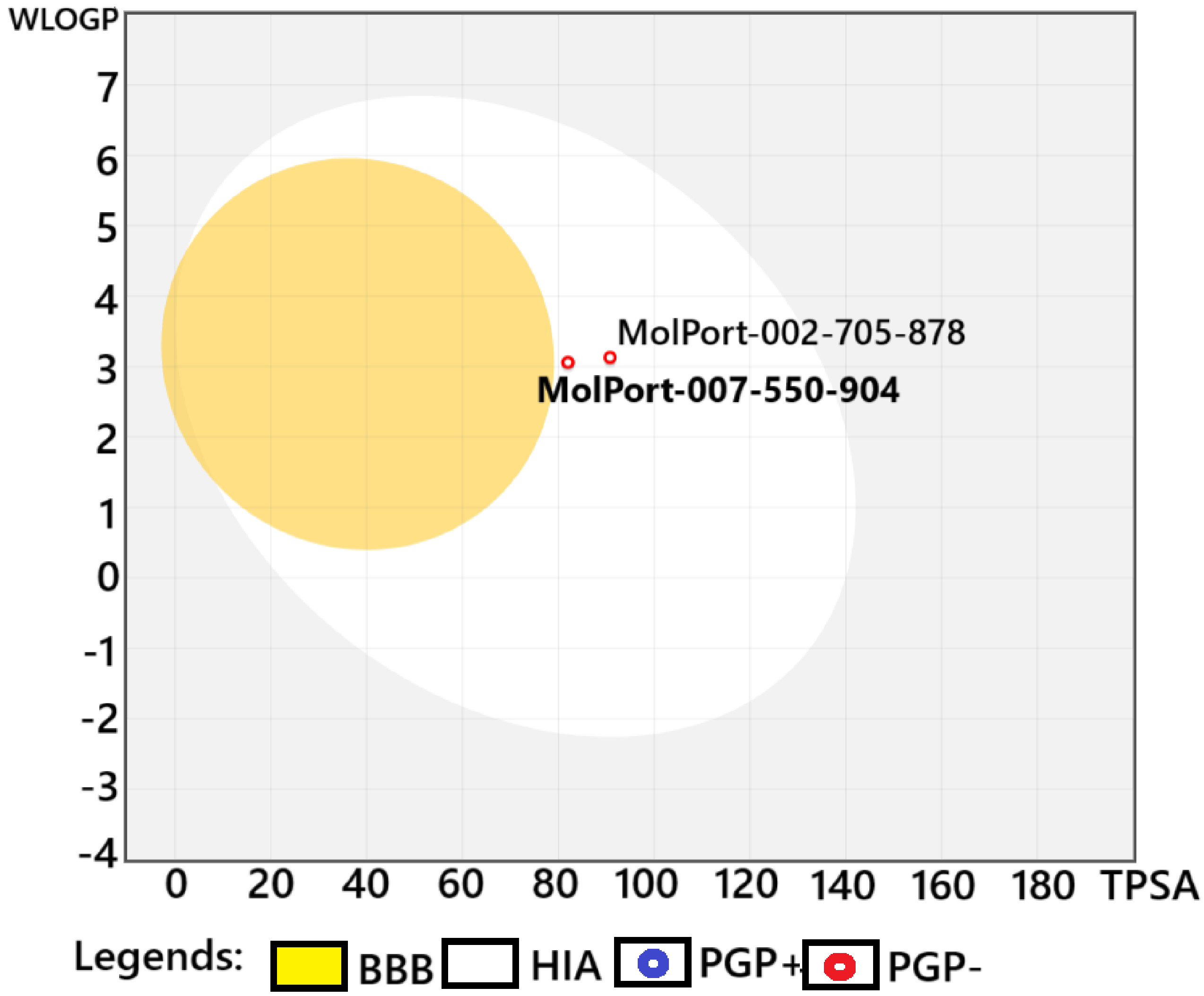


Figure 2S: Boiled-Egg image Molport-002-705-878 and Molport-007-550-904.
